# Unraveling SARS-CoV-2 Host-Response Heterogeneity through Longitudinal Molecular Subtyping

**DOI:** 10.1101/2024.11.22.624784

**Authors:** Kexin Wang, Yutong Nie, Cole Maguire, Caitlin Syphurs, Heejune Sheen, Meagan Karoly, Linda Lapp, Jeremy P. Gygi, Naresh Doni Jayavelu, Ravi K. Patel, Annmarie Hoch, IMPACC Network, David Corry, Farrah Kheradmand, Grace A. McComsey, Ana Fernandez-Sesma, Viviana Simon, Jordan P. Metcalf, Nelson I Agudelo Higuita, William B. Messer, Mark M. Davis, Kari C. Nadeau, Monica Kraft, Chris Bime, Joanna Schaenman, David Erle, Carolyn S. Calfee, Mark A. Atkinson, Scott C. Brackenridge, David A. Hafler, Albert Shaw, Adeeb Rahman, Catherine L. Hough, Linda N. Geng, Al Ozonoff, Elias K. Haddad, Elaine F. Reed, Harm van Bakel, Seunghee Kim-Schultz, Florian Krammer, Michael Wilson, Walter Eckalbar, Steven Bosinger, Charles R. Langelier, Rafick P. Sekaly, Ruth R. Montgomery, Holden T. Maecker, Harlan Krumholz, Esther Melamed, Hanno Steen, Bali Pulendran, Alison D. Augustine, Charles B. Cairns, Nadine Rouphael, Patrice M. Becker, Slim Fourati, Casey P. Shannon, Kinga K. Smolen, Bjoern Peters, Steven H. Kleinstein, Ofer Levy, Matthew C. Altman, Akiko Iwasaki, Joann Diray-Arce, Lauren I. R. Ehrlich, Leying Guan

**Author notes:** IMPACC Network group members (listed in Pubmed) are listed at the end of the document. Co-first authors (Kexin Wang, Yutong Nie, Cole Maguire).

## Abstract

Hospitalized COVID-19 patients exhibit diverse immune responses during acute infection, which are associated with a wide range of clinical outcomes. However, understanding these immune heterogeneities and their links to various clinical complications, especially long COVID, remains a challenge. In this study, we performed unsupervised subtyping of longitudinal multi-omics immunophenotyping in over 1,000 hospitalized patients, identifying two critical subtypes linked to mortality or mechanical ventilation with prolonged hospital stay and three severe subtypes associated with timely acute recovery. We confirmed that unresolved systemic inflammation and T-cell dysfunctions were hallmarks of increased severity and further distinguished patients with similar acute respiratory severity by their distinct immune profiles, which correlated with differences in demographic and clinical complications. Notably, one critical subtype (SubF) was uniquely characterized by early excessive inflammation, insufficient anticoagulation, and fatty acid dysregulation, alongside higher incidences of hematologic, cardiac, and renal complications, and an elevated risk of long COVID. Among the severe subtypes, significant differences in viral clearance and early antiviral responses were observed, with one subtype (SubC) showing strong early T-cell cytotoxicity but a poor humoral response, slower viral clearance, and greater risks of chronic organ dysfunction and long COVID. These findings provide crucial insights into the complex and context-dependent nature of COVID-19 immune responses, highlighting the importance of personalized therapeutic strategies to improve both acute and long-term outcomes.

## Introduction

The coronavirus disease 2019 (COVID-19) pandemic has resulted in millions of hospitalizations and deaths globally^1^. Hospitalized COVID-19 patients exhibit a wide range of acute disease severities, from less severe illness to critical conditions involving organ damage and mortality ^2^. Beyond the acute phase, 40-80% of hospitalized COVID-19 patients develop post-acute sequelae of SARS-CoV-2 (PASC, also known as Long COVID), a complex condition characterized by heterogeneous and persistent symptoms that can last for months to years after the acute infection^3–5^.

Immune signatures associated with poor COVID-19 outcomes have been identified at both the acute and convalescent stages ^6–12^. Hallmarks of acute disease severity, measured by respiratory status and mortality, include over-production of proinflammatory cytokines^8,13,14^, lymphopenia^8,15–17^, formation of neutrophil extracellular traps (NETs)^18^, dysregulation of complement and coagulation^8,19,20^, impaired interferon (IFN) signaling^7,21–23^, immune senescence^24^, delayed onset of neutralizing antibodies^25^, and signatures of apoptosis and tissue damage^17,26^, which have been confirmed in our prior studies focusing on an inpatient cohort exclusively^6,7^. Nevertheless, despite the observed association between immune responses and disease severity, significant heterogeneity remains among patients with similar levels of acute illness.

Subtyping analysis of COVID-19 patients, particularly those with severe complications, provides critical insights into the variability of immune responses and their associations with acute and long-term clinical outcomes. However, studies in this area have been constrained by small cohorts, limited molecular assays, and short-term longitudinal data ^27–30^. Here, we address these limitations by constructing patient subtypes using large-scale, longitudinal multi-omics data—including transcriptomics, proteomics, and metabolomics—collected from over 1,000 hospitalized patients in the IMPACC (IMmunoPhenotyping Assessment in a COVID-19 Cohort) study during acute infection (the first 28 days post-hospital admission), followed by PASC symptom survey monitored for up to one-year post-discharge (Figure 1A-B). We identified five subtypes with distinct immune trajectories, highlighting their heterogeneity in immune responses and links to pre-existing conditions, acute clinical outcomes, and post-acute recovery in a unified framework (Figure 1C).

**Figure 1:**
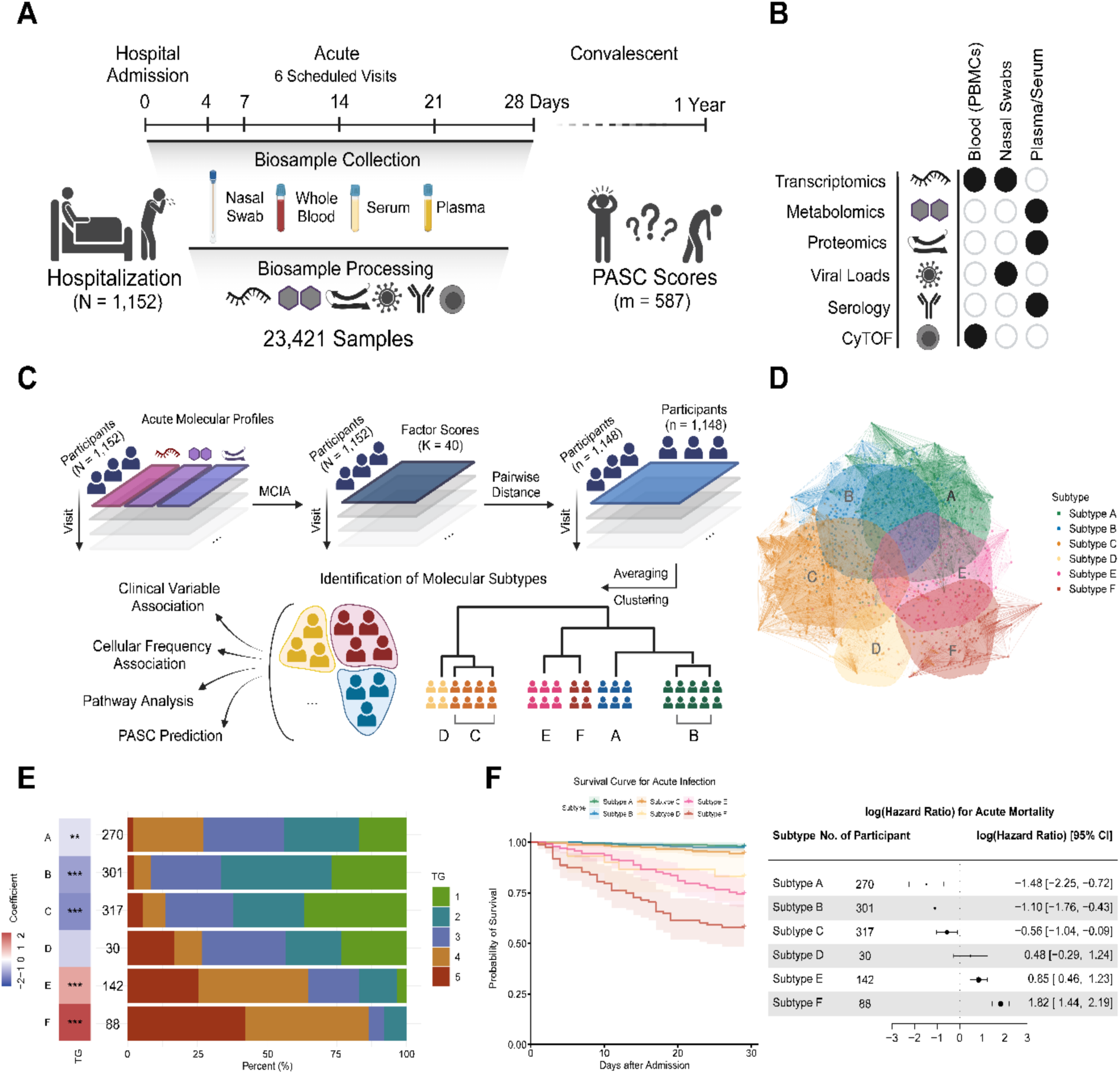
Analysis pipeline and acute primary clinical outcomes. **(A)** Timeline showing study design, patient hospitalization (N=1,152), biosample collection and processing (n=23,421), and PASC group (m=587) based on self-reported PASC surveys monitored up to one-year post-discharge. **(B)** Measured omics assays across various biosamples, including transcriptomics, metabolomics, proteomics, viral loads, serology and CyTOF. **(C)** Pipeline for constructing disease subtypes among patients using multi-omics molecular profiles and downstream analysis. **(D)** Network plot illustrating the multi-dimensional scaling of the distance between participants. Each ellipse encompasses 60% of the points from the centroid of its respective cluster. **(E)** Comparisons of acute severity level across subtypes using ordinal regression test and TG group assignment with TG1 being the least severe and TG5 being the most severe, with barplot presenting the empirical TG proportions within each subtype and its row legend presenting the ordinal regression coefficients and significance levels (*p<=0.05, **p<=0.01, ***p<=0.001). **(F)** Survival curve and forest plot showing the log-transformed hazard ratios for acute mortality for each subtype. In the left panel, shaded areas indicate the 95% confidence intervals from Kaplan-Meier estimates (thick line). In the right panel, horizontal lines represent the 95% confidence intervals for log hazard ratios from Cox proportional-hazards model controlling for age and sex. Intervals covering 0 suggest non-significant differences compared to other subtypes.

## Results

### Overview of the IMPACC Cohort and Analysis Pipeline

A total of 1,152 vaccine-naive participants hospitalized with COVID-19 were enrolled in the IMPACC study across 20 US hospitals between May 2020 and March 2021. Deep immunophenotyping was conducted at six visits in the first month of hospitalization^6,7,31^ (Figure 1A, Extended Data 1A). A total of 23,421 immunophenotyping samples, including nasal and PBMC transcriptomics, plasma metabolomics, plasma and serum proteomics, whole blood CyTOF, nasal viral load, and serology, were collected and analyzed to examine host immune response heterogeneity comprehensively (Figure 1B). Participants during the acute infection phase were categorized into five acute clinical trajectory groups based on modified WHO scores as described in our prior work^2^. From most mild to fatal, these were: Trajectory Group 1 (TG1, n=255), Trajectory Group 2 (TG2, n=306), Trajectory Group 3 (TG3, n=272), Trajectory Group 4 (TG4, n=212), and Trajectory Group 5 (TG5, n=107). Most TG1-3 patients were discharged within two weeks, while TG4-5 patients typically required mechanical ventilation, had prolonged hospital stays (≥28 days, TG4), or experienced fatal outcomes by Day 28 (TG5)^2^. Of these participants, 587 completed quarterly patient-reported outcome surveys up to one-year post-hospital discharge (Figure 1A), allowing classification into four groups: minimal deficit (n=356), physical deficit (n=92), cognitive deficit (n=81), and multiple deficits (n=58)^32^, with the latter three considered evidence of PASC. See Table S1 for detailed patient characteristics.

We used multiple co-inertia analysis (MCIA) to derive low-dimensional multi-omics factors from 17,569 acute molecular features covering transcriptomics, proteomics, and metabolomics^33^. These multi-omics factors captured major covarying patterns in high-dimensional molecular features across blood and nasal samples and were used to calculate pairwise distances among 1,148 participants with available data from visit 1, incorporating multiple visit samples where possible. Hierarchical clustering of these distances identified subtypes (referred to as molecular subtypes to reflect its dependence on the molecular profiles) separating 1,148 participants. We associated these subtypes with demographic and clinical characteristics to explore whether heterogeneous immune responses were linked to clinically relevant outcomes (Figure 1C).

We hypothesized that these unbiased subtypes would reveal immune heterogeneity beyond what is captured by acute respiratory status and mortality. To further characterize these subtypes, we evaluated their molecular functions and examined correlations with viral loads, antibody titers, and CyTOF cell frequencies. Additionally, we developed machine learning models to predict PASC development using acute molecular profiles and subtypes, comparing them to models based solely on clinical data to assess whether early molecular features offered added predictive value for PASC outcomes.

### Molecular Subtypes Exhibit Distinct Acute Clinical Characteristics

We identified six molecular subtypes, denoted as SubA, SubB, SubC, SubD, SubE and SubF, using hierarchical clustering of multi-omics factors during acute COVID-19 infection (Figure 1D, Extended Data1B-F and Table S1). Compared to SubA-SubC (n=270, 301, and 317, respectively), SubE and SubF (Figure 1E, n=142 and 88, respectively) were associated with more acute clinical trajectory groups TG4 and TG5 (p.val = 1.3E-12 and 2.7E-30). SubF also exhibited a significantly higher risk of mortality during the acute phase compared to all other subtypes (Figure 1F, p.val = 2.3E-21), followed by SubE (p.val = 1.6E-5). Furthermore, SubF had a significantly higher acute mortality rate than SubE (Extended Data 1F, p.val = 1.8E-4). SubD (n=30) had a mixed acute severity profile and could potentially be interesting for further investigation, but due to its small sample size, we restricted our focus to the three severe subtypes SubA-SubC and two critical subtypes SubE-SubF during acute infection.

We next examined whether the subtypes were associated with different demographics and comorbidities (Table S2). Among the critical subtypes, SubF had the highest acute mortality but a lower average admission age than SubE, which had the highest admission age (Figure 2A-B, adj.p = 1.2E-11). SubC exhibited a similar average admission age as SubF, followed by SubA and SubB which has the lowest age at admission (adj.p = 2.8E-13). Ethnic and gender differences were also observed: SubA had the highest proportion of Hispanic or Latino participants (Figure 2A, 2C, adj.p = 2.0E-7), and SubC had the highest proportion of females (adj.p = 1.8E-6). BMI was relatively comparable across subtypes with SubE demonstrating a moderately lower rate of obesity (adj.p = 2.0E-2). 5 out of 13 comorbidities were significantly different (Table S2), with SubA and SubB having considerably fewer comorbidities relative to other groups (Figure 2A, 2D).

**Figure 2:**
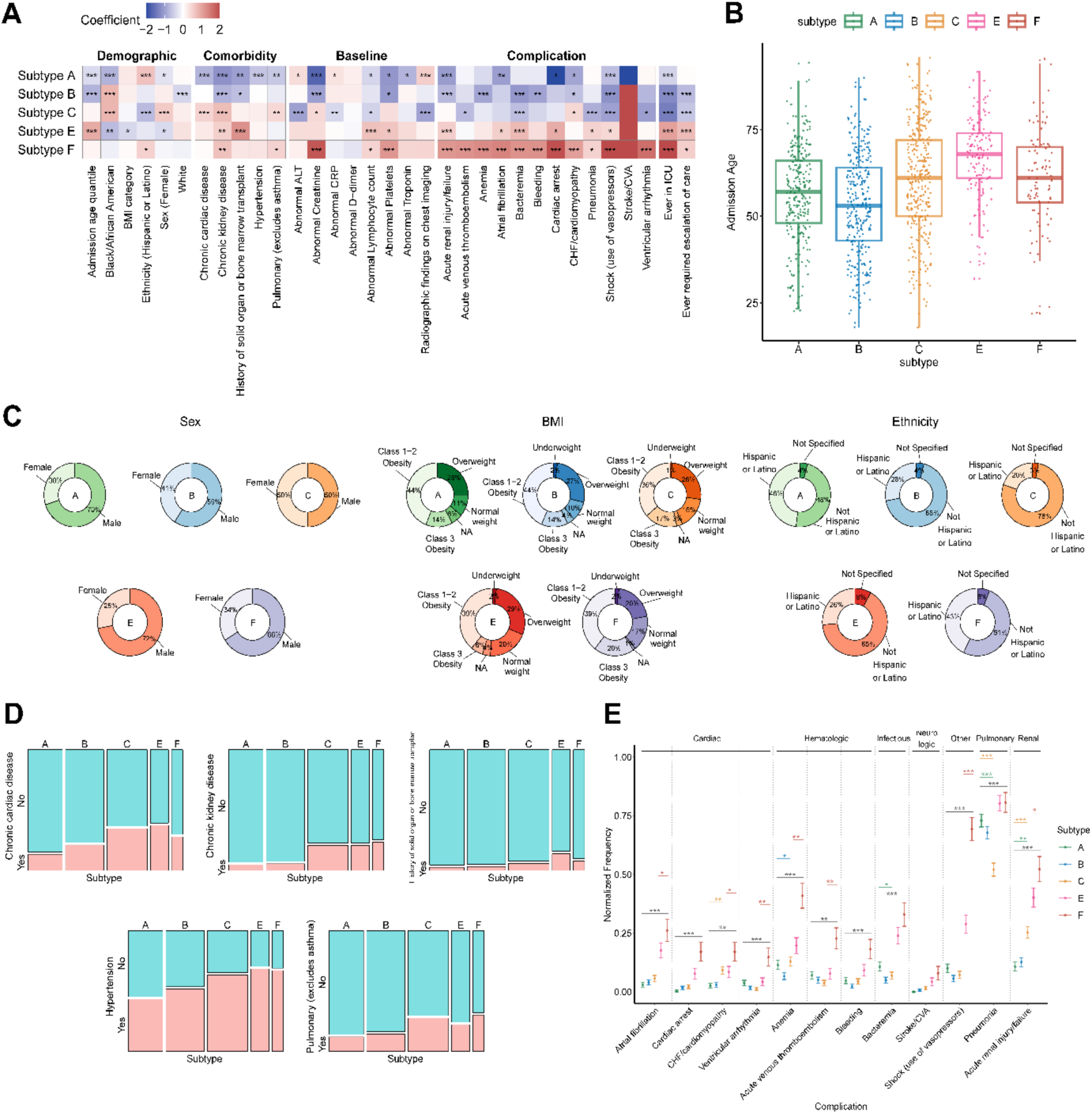
Patient molecular subtypes exhibit distinct acute clinical characteristics. **(A)** Heatmap of coefficients and BH adjusted p-values comparing clinical characteristics across subtypes, with clinical characteristics including demographics, comorbidities, baseline laboratory tests, complications and hospital course. Each subtype was compared against all others using generalized linear regressions (ordinal for admission age quantiles, BMI, and radiographic findings on chest imaging, and logistic regression for others). Except for demographics, models have been adjusted for age and sex. **(B)** Boxplot of admission age distribution. **(C)** Donut plots for subtype demographics, including sex, BMI, and ethnicity. **(D)** Bar plots of top differentially enriched comorbidities across subtypes (Chi-squared test adj.p<=0.001). **(E)** Normalized frequency plot showing top differentially enriched complications across subtypes (Chi-squared test adj.p<=.001) grouped by bodily systems and functions. Vertical bars indicate the mean frequency and the corresponding standard error. Horizontal lines marked with asterisks show the significance of group-wise comparisons, with the grey line indicating the comparison between EF and ABC. Lines and asterisks matching the colors of A, B, and C denote comparisons of each group against the other two – A vs BC, B vs AC, and C vs AB respectively. The red line matching the color of F represents the comparison between F and E. (*adj.p<=0.05, **adj.p<=0.01, ***adj.p<=0.001).

The five patient subtypes also exhibited distinct complications during hospitalization (Figure 2A, 2E). Subtypes SubE and SubF showed significantly higher rates of complications compared to SubA-C, consistent with previously reported associations between complications and acute clinical severity measured by respiratory status and mortality^2^. Notably, several differences existed even among subtypes with similar acute severity. Among the critical subtypes, SubF experienced higher rates of shock (adj.p = 1.1E-9), acute renal failure (adj.p = 3.7E-2), hematologic complications such as anemia and acute venous thromboembolism (adj.p = 1.8E-3 and 4.9E-3), and cardiac complications, including atrial fibrillation, congestive heart failure (CHF), and ventricular arrhythmia (adj.p = 4.0E-2, 2.8E-2, 1.0E-2). Within the severe SubA-C subtypes, SubC had the highest rates of CHF and acute renal failure (adj.p = 1.0E-3 and 2.5E-6), but a lower incidence of pneumonia (adj.p = 1.9E-6). These significant associations between subtypes and complications persisted even after adjusting for five significant comorbidities from the comorbidity analysis and demographics (Extended Data 2A). Similar patterns were observed after grouping complications by bodily systems and functions (Extended Data 2B-C).

### Cellular Compositions and Serum Proteins Differentiated Subtypes

Serum soluble proteins were clustered into ten modules (SPmod1-SPmod10) using hierarchical clustering, revealing significant differences across patient subtypes (Extended Data 3A-B). In this section, we investigated the key serum protein profiles distinguishing these subtypes, along with differences in whole blood cellular composition, nasal viral load, and SARS-CoV-2-specific immunoglobulin (Ig) levels (Extended Data 3C).

Critical subtypes (SubE and SubF) exhibited slightly higher nasal viral loads compared to averaged severe subtypes that were not statistically significant (Figure 3A). However, SubE and SubF showed persistent elevations in neutrophil and monocyte frequencies, along with T-cell lymphopenia (Figure 3A-B, adj.p < 0.05, and Extended Data 3D), which are consistent with alterations in circulating blood cell composition reported in severe/critical COVID-19 cases^6–8^. SubE and SubF also showed persistent elevation in many cytokines, such as IL-6 and IL-10, and were particularly outstanding for serum proteins in SPmod1/2/7/9, which are involved in immune regulation and inflammation (Figure 3D-E, adj.p < 0.05). These elevations reflected a hyperinflammatory state, consistent with uncontrolled inflammatory cytokine production reported in severe/critical patients^8,13,14,34^. Additionally, SubE and SubF showed reduced levels of proteins in SPmod4 which included FLT3LG, TNFSF11 (RANKL), and IL12B (p40) that are integral to T-cell production, activation, and function. Transcriptomic signatures for T-cell cytotoxicity were also reduced in SubE and SubF (Extended Data 3E).

**Figure 3:**
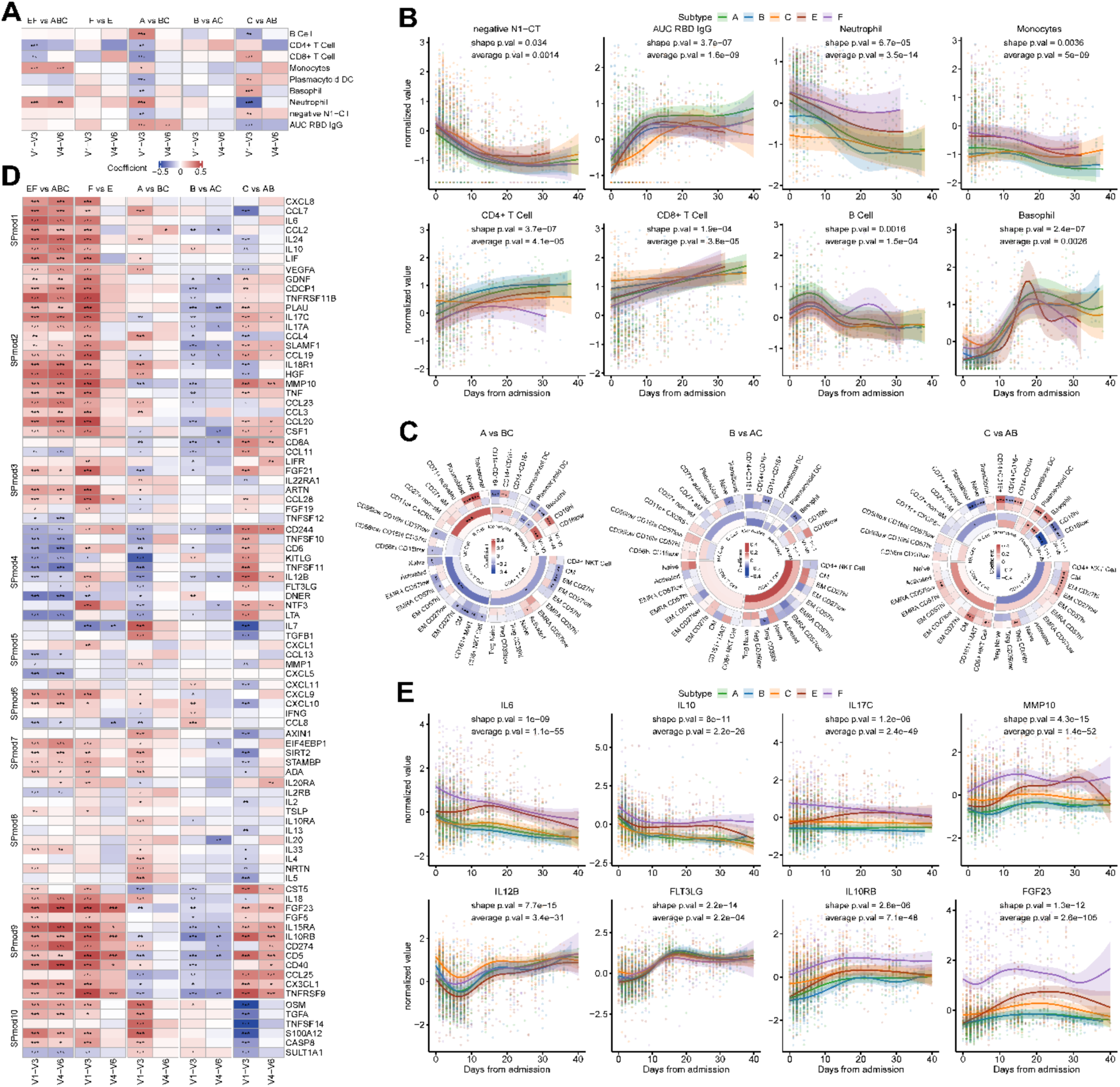
Molecular subtypes exhibit distinct cell composition and serum protein characterization. **(A)** Heatmap of comparisons in cell composition, viral load and antibody for visits 1-3 (less than 10 days after admission) and visits 4-6 (more than 10 days after admission) among 5 major subtypes using linear mixed effect modeling after further adjusting for participant ID and enrollment site as random effects, and age, sex, and admission date as fixed effects, with heatmap cell colored based on regression coefficient (white if adj.p > 0.2) and annotated by its significance. **(B)** Longitudinal trajectories of negative N1-CT, AUC RBD IgG, neutrophil, monocytes, CD4+ T cell, CD8+ T cell, B cell and basophil, colored by subtypes. Shaded region denotes 95% confidence interval from generalized additive mixed model of the fitted trajectory (thick line). **(C)** Circular heatmap of comparisons in whole blood (CyTOF) of both parent and child populations for visits 1-3 and visits 4-6 among severe subtypes. **(D)** Heatmaps of comparisons in serum protein profiles for visits 1-3 and visits 4-6 among 5 major subtypes, with proteins grouped based on SPmod and heatmap cells colored by the regression coefficients (white if adj.p > 0.2). **(E)** Longitudinal trajectories of a small subset of most notable serum proteins IL-6, IL-10, IL17C, MMP10, IL12B, FLT3LG, CD274 and FGF23, colored by subtypes. Shaded region denotes 95% confidence interval from generalized additive mixed model of the fitted trajectory (thick line). (*adj.p<=0.05, **adj.p<=0.01, ***adj.p<=0.001).

Among the severe subtypes, SubA exhibited the lowest, and SubC had the highest viral loads at early visits (adj.p = 1.3E-3/4.0E-3, Figure 3A-B). These differences in viral loads were accompanied by distinct antiviral responses. SubA demonstrated a controlled early increase in innate immune responses alongside a strong humoral response, while SubC was characterized by robust early T-cell immunity but insufficient humoral response. SubB, positioned between SubA and SubC, exhibited a more balanced cellular response. Specifically, SubA had relatively higher frequencies of neutrophils, monocytes, and B cells, but lower counts of CD4+ and CD8+ T cells at the first three visits (adj.p: neutrophils = 4.9E-7, monocytes = 2.1E-2, B cells = 2.0E-6, CD4+ T cells = 3.6E-3, CD8+ T cells = 1.4E-5). Conversely, SubC exhibited the highest frequencies of CD4+ and CD8+ T cells but the lowest neutrophil, monocyte, and B-cell frequencies at early visits (adj.p: neutrophils = 6.0E-11, monocytes = 0.108, B cells = 1.5E-3, CD4+ T cells = 0.138, CD8+ T cells = 6.4E-5), as well as the lowest baseline absolute neutrophil count (Extended Data 3D). The elevated CD8+ T-cell frequencies in SubC suggested enhanced cytotoxic T-cell responses, which were supported by its elevated transcriptomic signatures for T-cell cytotoxic functions at early visits (Extended Data 3E). The early differences diminished over time. Further analysis of cell sub-populations revealed that classical CD14+CD16-monocytes (adj.p = 1.7E-3/2.0E-4 for SubA vs SubB-C and SubC vs SubA-B respectively), naive B cells (adj.p = 4.5E-10/7.4E-5), plasmablasts (adj.p = 2.4E-7/2.7E-10), CD4+ T_CM/EM_ cells (adj.p: CM = 1.2E-3/6.5E-5, EM CD27hi = 2.9E-5/7.2E-5), and CD8+ T_CM_ cells (adj.p = 2.2E-4/9.2E-9) were significant contributors to the observed differences in the parent cell populations among the severe subtypes (Figure 3C, Extended Data 4A-B). Additionally, SARS-CoV-2-specific IgG levels were persistently highest in SubA, especially during the early visits (adj.p = 4.9E-7 at early visits; adj.p = 2.7E-3 at late visits) and lowest in SubC (adj.p = 4.5E-5 at early visits; adj.p = 5.4E-2 at late visits). Although sensitive to timing, the differences in viral loads and humoral responses across subtypes remained significant after adjusting for days from symptom onset (Extended Data 4C-D).

The cellular differences among SubA, SubB and SubC were also reflected in their distinct serum protein profiles (Figure 3D-E and Extended Data 4E). Consistent with SubA’s early increase in neutrophils and classical monocytes, SubA exhibited an early elevation in SPmod1, which includes immunomodulatory cytokines and chemokines such as IL-6, IL-10, CCL2, and CCL7. Notably, CCL7, a chemoattractant particularly active for monocytes, was highest in SubA and lowest in SubC. SPmod2 was lowest in SubB and showed mixed patterns in SubA and SubC, with MMP10 and CSF1 showing persistent elevation in SubC. Both MMP10 and CSF1 have been associated with COVID-19 severity, atherosclerosis, and increased carotid intima-media thickness^35,36^, aligning with the higher incidence of cardiac complications in SubC. CSF1 (M-CSF) is central to monocyte development and differentiation into tissue-resident macrophages^37^, with elevated CD16+ monocytes observed after M-CSF increase^38^, corroborating elevated CD16+ monocytes in SubC (Figure 3C, adj.p < 0.05).

Reflecting the higher T-cell presence in SubC versus SubA at early visits, SPmod4 (including FLT3LG, TNFSF11, IL12B) was broadly higher in SubC and lower in SubA early on. FLT3LG and TNFSF11 enhance T-cell function by promoting the proliferation and differentiation of dendritic cells^39–41^, potentially linking lower T-cell production to fewer pDCs (plasmacytoid dendritic cells) in SubA at early visits (Figure 3A, adj.p = 8.8E-4). Although SPmod5 showed less distinct patterns across SubA-C, IL7 was much higher in SubA and lower in SubC. IL-7 is crucial for T-cell survival, development, and homeostasis, and influences early B-cell development^42,43^. Further, SubC showed elevated levels of SPmod9 compared to SubA and SubB (Figure 3D-E, Extended Data 4E). SPmod9 includes immune-regulating proteins such as TNFRS9 (41BB), which enhances T-cell function and has been linked to chronic inflammatory condition^44^, and IL-10RB, a shared subunit for IL-22 and IL-26 receptor complexes with both cytokines primarily produced by T cells and associated with chronic immune disorders^45,46^. SPmod9 also includes FGF5 and FGF23, with FGF23 strongly associated with chronic kidney disease^47,48^, potentially explaining SubC’s higher incidence of cardiovascular and renal complications compared to SubA and SubB.

Although SubE and SubF did not exhibit major differences in circulating cellular compositions (Figure 3A-B), SubF displayed a broader elevation of inflammatory serum proteins relative to SubE during early visits (Figure 3D-E), indicating a more severe hyperinflammatory state, which aligns with the increased incidence of complications in SubF.

### Multi-omic Factors of Blood and Upper-airway Inflammation Distinguish Patient Subgroups

The subtypes were constructed based on the pairwise patient distances of their longitudinal multi-omics factors, with each factor capturing strong covarying patterns among observed high-dimensional analytes. To further understand the immune programs underpinning these subtypes, we selected multi-omics Factors 1, 2, 3, and 10 as the top discriminating factors separating them for further investigations (Supplementary Materials).

Factor 1 showed persistent elevation in critical subtypes SubE-F relative to severe subtypes SubA-C, while being elevated in SubA-B relative to SubC only in early visits (Figure 4A). Functionally, Factor 1 captures a state of coordinated and widespread immune dysregulation and contains many high-contributing analytes, defined as those strongly associated with a factor, from different assays across blood and nasal compartments. Specifically, Factor 1 featured the top 30 enriched pathways indicative of systemic inflammation (NET formation = 9.4E-10; complement and coagulation adj.p = 1.1E-9; Inflammatory response adj.p = 3.5E-6, ROS adj.p=4.1E-6) and suppressed T cell functions (Th1 and Th2 differentiation adj.p = 2.1E-9, T-cell receptor signaling adj.p = 2.2E-9, Th17 differentiation adj.p = 2.2E-9) (Figure 4B), which echoed the “severity factor” associated with increased acute disease severity in the IMPACC cohort in our prior study^7^. Factor 1 also captured suppressed plasmalogen (adj.p=3.7E-6, including many plasmenyl-PC metabolites) and sphingomyelins (adj.p = 1.6E-5, including sphingomyelin with various chain lengths). Plasmenyl-PC is known for its roles in membrane fluidity and antioxidative properties, with lower levels associated with the occurrence and severity of COVID-19^49–51^. Additionally, SARS-CoV-2 virus activates acid sphingomyelinase as a crucial step of infection, and a low level of sphingomyelin has been associated with severe COVID^52,53^. These findings, along with elevated myeloid cells and inflammatory cytokines (e.g., IL-6, IL-10 from SPmod1) and reduced T-cells and related serum protein activities (SPmod4), indicate that unresolved systemic inflammation and T-cell dysfunction are key drivers of severity in SubE and SubF.

**Figure 4:**
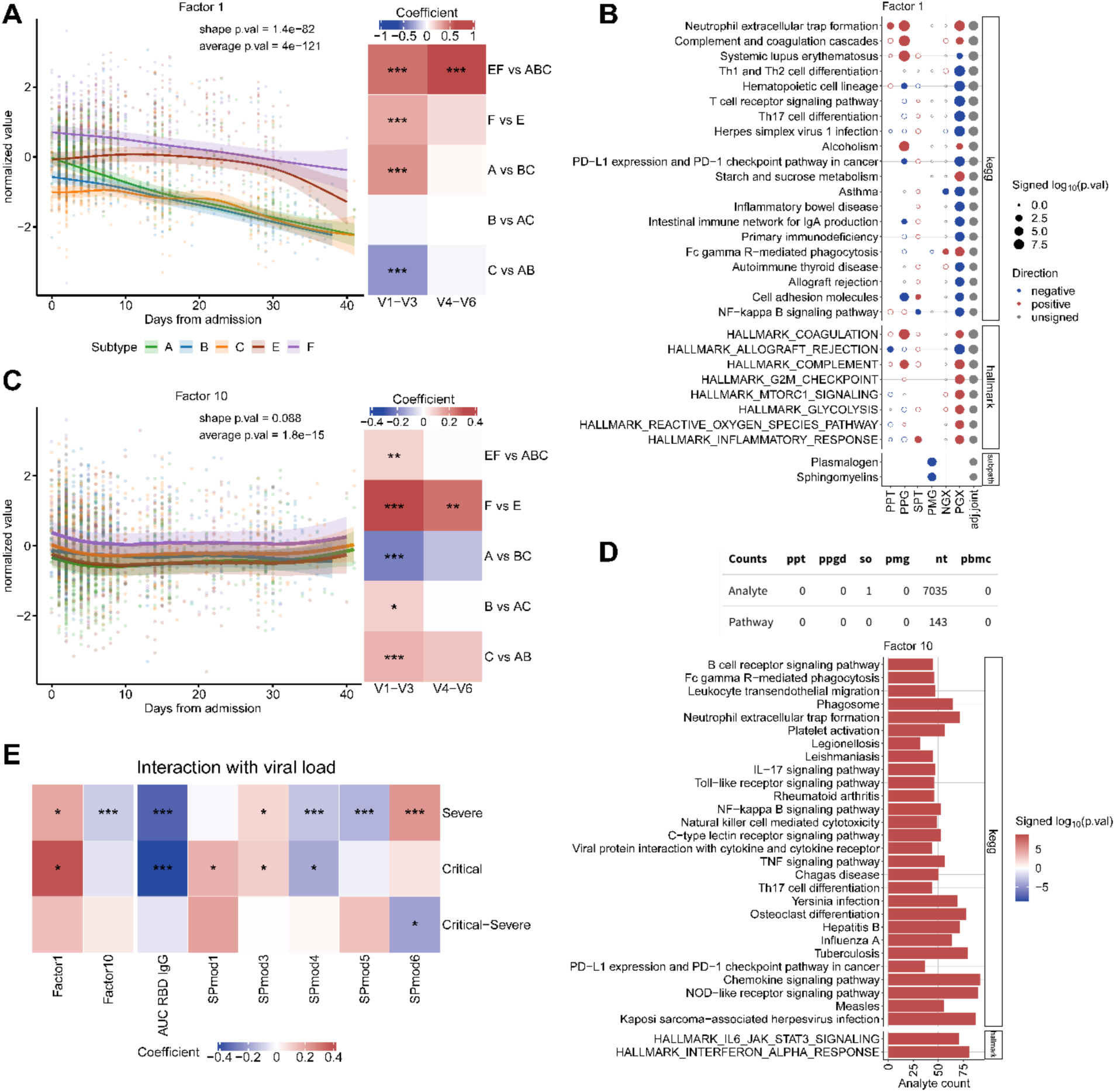
Blood and upper-airway inflammation underpins subtype heterogeneity. **(A)** Longitudinal trajectories of Factor 1 (right), colored by subtypes and with shaded region denoting 95% confidence interval, and heatmap of comparisons in Factor 1 for visits 1-3 (<10 days after hospitalization) and visits 4-6 among subtypes (left), colored by regression coefficients from the linear mixed effect model. **(B)** Pathway Enrichment of Factor 1, where column names PPT/PPG=Plasma Proteomics Targeted/Global, SPT =Serum Proteomics Targeted, PMG = Plasma Metabolomics Global, NGX/PGX=Nasal/PBMC gene expression, adj.joint = aggregated p-value across omics after BH correction. For Factor 1, only top 30 pathways measured by adj,joint out of 99 all pathways enriched with adj.joint < 0.05. **(C)** Longitudinal trajectories of Factor 10 and heatmaps of comparisons in Factor 10 for visits 1-3 and visits 4-6 among subtypes. **(D)** Pathway enrichment of Factor 10 (adj.joint < 0.05). The top table showed the high-contributing features and number of all pathways with unadjusted joint p.value < 0.05, which were all from NGX. **(E)** Regression coefficients from linear mixed effect model between nasal viral load and Factor 1, Factor 10, antibody and significant serum protein modules, for severe subtypes (SubA & B & C), critical subtypes (SubE & F), and the difference between critical and severe subtypes displaying interactions of critical subtypes compared to severe subtypes, respectively. The results for multi-omics factors Factor 1/10 were further adjusted for the more specific immune components on the right panels. (*adj.p<=0.05, **adj.p<=0.01, ***adj.p<=0.001).

On the other hand, Factor 10 was lower in the severe SubA and critical SubE, with SubC being the most elevated among severe subtypes (Figure 4C). It was primarily driven by nasal transcriptomics and indicated upper-airway pan-inflammation (Figure 4D). 7035 out of 7036 total high-contributing analytes for Factor 10 were nasal genes, whose top enriched pathways includes JAK-STAT3 signaling (adj.p = 2.0E-9, including innate Immune activators and inflammation mediator MYD88, TLR2, TNFRSF1A, IL1B, IL17RA,IL4R, TLR2, STAT3, TNF, IL-6, IFNGR1 and others), NET formation (adj.p = 2.8E-9), interferon alpha response (adj.p = 2.3E-9, including genes integral to the cellular response and regulatory mechanisms triggered by IFNa such as TRIM25, IL4R, and IRF1) and BCR signaling (adj.p = 1.9E-9, including key components in the early stages of BCR signaling such as GRB2, LYN, SYK, PLCG2, BTK, and downstream of BCR signaling such as NFKBIA, FOS). While elevated Factor 10 in SubF was compatible with its consistently high systemic inflammatory signatures, even compared to SubE at early visits, the elevation of Factor 10 in SubC was interesting given the subtype’s less severe systemic inflammation.

To further investigate the immune dynamics across subtypes, we analyzed the associations between viral load and Factors 1 and 10, anti-RBD titers, and SPmods using mixed-effects regressions (Figure 4E). Negative associations between immune profiles and viral load were interpreted as indicators of effective viral control. Positive associations between immune profiles and viral load were interpreted as viral-driven overproduction of immune responses. Among both severe and critical subtypes, SARS-CoV-2-specific IgG emerged as the strongest associate of lower viral control (adj.p = 1.0E-24/2.6E-14). Additionally, SPmod4, which is associated with enhanced T-cell frequencies and cytotoxic activity, displayed significant negative correlations with viral load in both groups (adj.p = 1.5E-3/4.5E-2), emphasizing the critical role of T-cell-mediated responses in controlling viral replication. We also identified several context-dependent immune responses. SPmod1, which includes key analytes linked to systemic inflammation and COVID-19 severity (e.g., IL-6, IL-10), demonstrated a significant positive correlation with viral load in the critical subtypes SubE and SubF (adj.p = 4.5E-2). This suggested that excessive inflammation in critical subtypes contributed to worsened outcomes in an immune environment already characterized by severe dysregulation. Indeed, SubF exhibited both higher levels of SPmod1 and a broader range of complications compared to SubE but SubA-B exhibited no more complications compared to SubC, suggesting a pathogenic role of uncontrolled inflammation in the critical subtypes but a salutary effect of early controlled inflammation among the less severe patients. SPmod6, which includes the chemokines CXCL9/CXCL10/CXCL11 that play critical roles in recruiting T-cells to sites of inflammation, showed a unique positive association with viral load in the severe subtypes that exhibited more abundant T-cell immunity (adj.p = 5.7E-10), implying a virus-driven T-cell site recruitment. Interestingly, Factor 10, the nasal inflammation factor, was a significant negative predictor of viral load only in the severe subtypes. This finding implies that the elevated upper-airway inflammation observed in SubB and SubC, relative to SubA, may act as a compensatory mechanism, aiding in viral control despite their lower humoral responses.

### SubF is Marked by Insufficient Anticoagulation and Fatty Acid Disruptions

Factor 2 showed a unique pattern differentiating among the critical subtypes and among the severe subtypes, with lower levels in SubF relative to SubE and in SubC relative to SubA-B (Figure 5A). This factor was functionally enriched in complement and coagulation-related plasma proteins (e.g., complement and coagulation (CC) cascade, adj.p = 4.2E-3), was associated with the decrease of a large family of fatty acids (FAs), including long-chain polyunsaturated FAs (PUFAs), long-chain monounsaturated FAs (MUFAs), long-chain saturated fatty acids (SFAs), and their derivatives (e.g., endocannabinoids, monohydroxy FAs, and branched FAs, adj.p = 9.4E-5/3.0E-4/2.9E-2, respectively).

**Figure 5:**
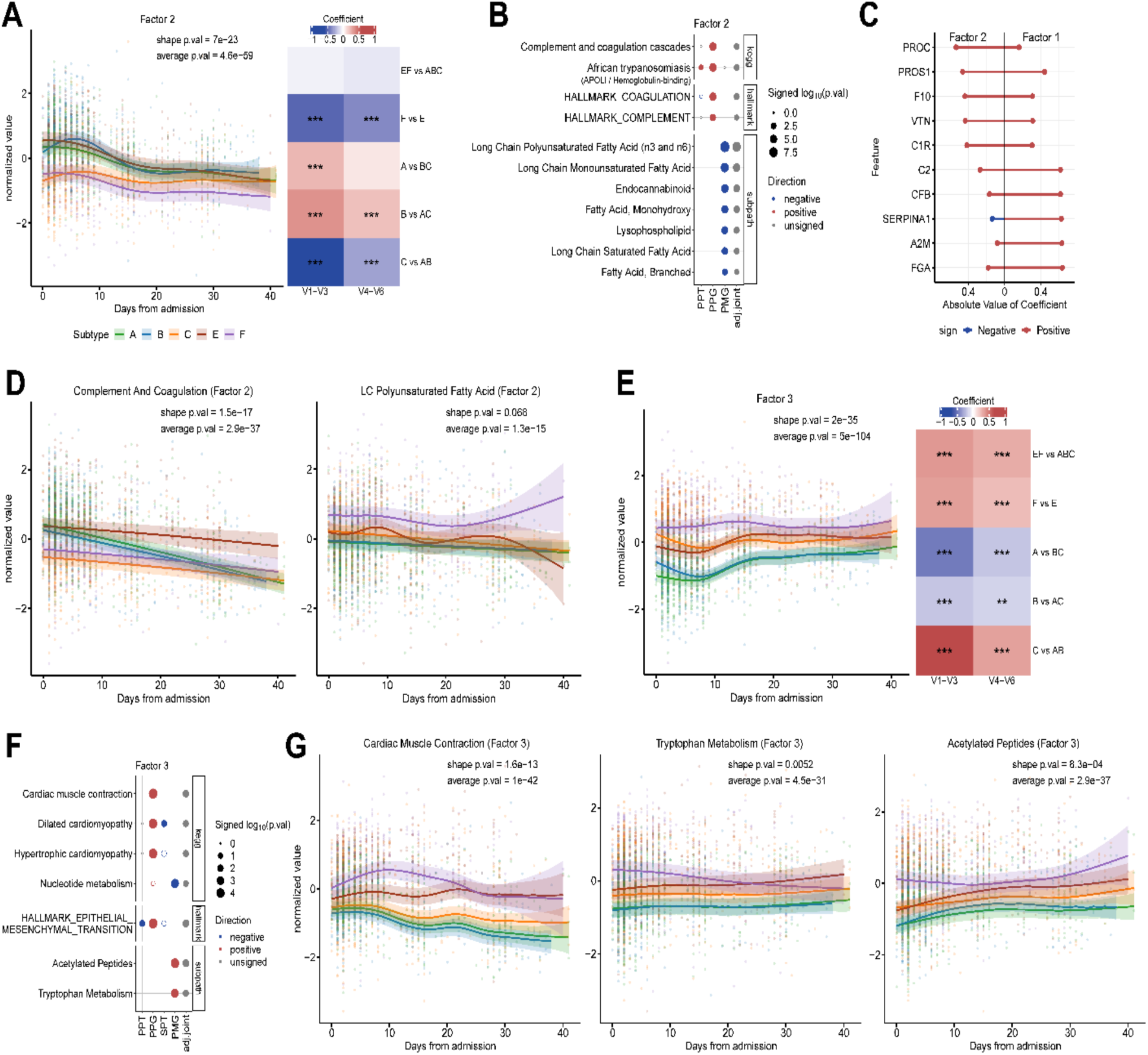
Dysregulated coagulation, disrupted fatty acid metabolism, and increased cardiomyopathy and amino acid catabolism identify COVID subtypes with increased complications. **(A)** Longitudinal trajectories of Factor 2 (left), colored by subtypes with shaded region denoting 95% confidence interval, and heatmaps of comparisons in Factor 2 for visits 1-3 and visits 4-6 among subtypes (right), colored by regression coefficients from the linear mixed effect modeling. **(B)** Pathway Enrichment of Factor 2 (adj.joint < 0.05). **(C)** Coefficient comparison for proteins enriched in complement and coagulation in Factor 1 and Factor 2. **(D)** Longitudinal trajectories of complement and coagulation and polyunsaturated fatty acid indicated in Factor 2, colored by subtypes. **(E)** Longitudinal trajectories of Factor 3, colored by subtypes, and heatmaps of comparisons in Factor 2 for visits 1-3 and visits 4-6 among subtypes. **(F)** Pathway Enrichment of Factor 3 (adj.joint < 0.05). **(G)** Longitudinal trajectories of cardiac muscle contraction, tryptophan metabolism, and acetylated peptides indicated in Factor 3, colored by subtypes. (*adj.p<=0.05, **adj.p<=0.01, ***adj.p<=0.001).

While both Factor 1 and Factor 2 showed CC cascade enrichment, they captured opposing functions. Factor 1 captured CC cascade enrichment in both PBMC transcriptomics and plasma proteomics, involving key proteins like FGA, A2M, SERPINA1, CFB, and C2 (Figure 5C). These proteins play crucial roles in inflammation and coagulation and have been linked to hyperinflammation, hypercoagulation, and thrombosis risk^54–56^. In contrast, Factor 2’s CC analytes were predominant in plasma proteomics and included leading-edge proteins such as PROC, PROS1, F10, VTN, and C1R (Figure 5C). PROC is known for controlling microvascular thrombosis and inflammation, with PROS1 acting as its cofactor^57–59^. VTN regulates complement activation and may protect against complement-mediated damage^60,61^. The CC pathway activity in Factor 2 was higher in SubA-B than SubC during early visits, suggesting successful anticoagulation regulation in SubA-B alongside early elevation in Factor 1, which later resolved and became comparable across SubA-C (Figure 5D). Additionally, CC pathway activity in Factor 2 remained elevated in SubE but not SubF throughout the study period, despite persistently high Factor 1 levels in both subtypes, suggesting a uniquely severe CC imbalance in critical SubF. This imbalance aligned with SubF’s increased incidence of CC-related complications including acute venous thromboembolism and bleeding (Figure 2E), and its high abnormal platelet counts in baseline lab tests (Figure 2A).

Factor 2 also captured broad elevations and disruptions of various FAs in SubF (Figure 5B). This was confirmed by elevated levels of long-chain PUFAs (n3 and n6) in SubF compared to other subtypes (Figure 5D). While long-chain SFAs like stearate (18:0), myristate (14:0), palmitate (16:0), arachidate (20:0), and margarate (17:0) are generally considered pro-inflammatory^62,63^, with palmitate previously linked to adverse COVID-19 outcomes^64,65^, the elevation of PUFAs may reflect increased oxidative stress in SubF due to inflammation^66,67^. Indeed, both n3 and n6 PUFAs, including arachidonic acid (AA) and linoleic acid (LA) which have been observed to increase with COVID-19 severity^68^, were positively correlated with TNF at visit 1 (Extended Data 5A-B), and the n6/n3 ratios were predominantly elevated in SubF (Extended Data 5C), suggesting early systemic hyperinflammation^66^. The elevated oxidative stress hypothesis in SubF was consistent with SubF’s higher rate of anemia (Figure 2A and 2E), and further supported by increased levels of peroxidation products 13-HODE and 9-HODE^69^, which were leading-edge metabolites of the monohydroxy FAs pathway. Additionally, the FA disruptions in SubF were corroborated by reduced levels of APOLI/Hemoglobin-binding proteins in the plasma (Figure 5B, adj.p = 8.4E-3, including HBB, HBA2, HPR, SELE, APOL1, and APOA1 as leading-edge proteins). Lower levels of SELE, APOL1, and APOA1 may indicate alterations in lipid metabolism^70^, while reduced levels of HPR, HBB, and HBA2 could result from hemolysis due to high oxidative stress^71^. Moreover, altered levels of FAs have been associated with changes in the proteome, with CC pathway alterations most strongly linked to FA changes^66,72^.

### SubA and SubB Exhibited Reduced Cardiomyopathy Markers and Amino Acid Catabolism

Factor 3 distinguished the critical subtypes from the severe and was also more elevated in SubC compared to SubA-B (Figure 5E). This factor was enriched in pathways associated with chronic organ damage, including epithelial-mesenchymal transition (EMT) (with learning-edge proteins TPM1-4, LOX, MMP14, and various collagen types such as COL6A3, COL4A2, COL1A1, adj.p = 5.3E-3) and cardiomyopathy-related pathways (cardiac muscle contraction, dilated/hypertrophic cardiomyopathy all with adj.p < 0.05, and including large shared leading-edge proteins TPM1-4, ACTC1, ACTB, and HRC) (Figure 5F). The EMT leading-edge proteins indicated a strong presence of EMT and fibrotic processes, which also correlated with the elevation of cardiomyopathy-related proteins^73,74^. We confirmed that cardiac muscle contraction pathway activity was highest in the critical subtypes while being significantly higher in SubC compared to SubA and SubB (Figure 5G). Lower Factor 3 scores in SubA and SubB corroborated with their fewer chronic cardiac complications, while SubF exhibited the highest rate of cardiac complications, followed by SubE and then SubC (Figure 2E).

Factor 3 was also enriched in acetylated peptides (e.g., 4-hydroxyphenylacetylglutamine, phenylacetylglutamate, and phenylacetylglutamine, adj.p = 6.0E-3) and tryptophan catabolism (e.g., C-glycosyltryptophan, 8-methoxykynurenate, indoleacetylglutamine, N-acetyltryptophan, anthranilate, N-acetylkynurenine, kynurenate, kynurenine, indolelactate, 3-indoxyl sulfate, indolebutyrate, adj.p = 1.2E-2) in plasma metabolites (Figure 5F). These pathways showed distinct patterns across subtypes: SubA and SubB had the lowest activity, SubC showed moderate activity, and SubE and SubF exhibited the highest activity (Figure 5G). Leading-edge metabolites in these pathways, such as kynurenine and 3-indoxyl sulfate, have been associated with both chronic and acute kidney disease^75–78^, consistent with the lowest rates of chronic kidney disease and renal injury complications observed in SubA and SubB. Elevated phenylacetylglutamine has also been linked to overall mortality and several chronic cardiovascular diseases^79–81^. Given that the pathways enriched in Factor 3 suggested a state associated with chronic organ damage, we investigated whether differences in Factor 3 across subtypes could be explained by comorbidities. While chronic kidney and cardiac comorbidities were indeed significantly associated with Factor 3 (Extended Data 5D), significant differences across subtypes remained even after adjusting for comorbidities (Extended Data 5E).

### Molecular Subtypes Associate with Late-Stage Recovery

Our study included 586 participants with long COVID survey responses that classified them into minimal deficit (well-recovered), physical deficit, cognitive deficit, and multiple deficits, of whom 574 were included in our severe or critical subtypes (Extended Data 6A). Although the clinically defined acute TG groups did not contribute to the PASC assignments in all available participants^32^ or within the defined subtypes (Extended Data 6B), the patient subtypes identified in our study associated with distinct PASC patterns (overall p.val = 6.4E-3, Figure 6A): SubA contained the least proportion of participants with PASC deficit (p.val = 3.4E-3), SubC was enriched for participants with PASC, mainly in physical and multiple deficits (p.val = 4.4E-3/9.7E-2 respectively), and SubF was also enriched in PASC participants, primarily from the cognitive deficit group (p.val = 1.7E-3). The incomplete PASC assignments in our cohort resulted from non-respondents to patient surveys after hospital discharge, potentially indicating poor post-acute recovery. For example, 47 deceased participants within 3 months after hospitalization lacked long COVID survey data and hence, did not have PASC assignments. The rate of convalescent survey non-respondents was significantly biased across subtypes as measured by its relative ratio to the minimal deficit (Extended Data 6C), being significantly higher in SubF and SubC (p.val = 1.1E-2 and 1.3E-3 respectively), while being lowest in SubA (Extended Data 6C), consistent with inferior documented post-acute recovery for SubC and SubF and best recovery for SubA. These differential patterns remained consistent after adjusting for age and sex (Extended Data 6D). Moreover, SubF also exhibited a significant increase in mortality during the post-acute stage (Figure 6B, p.val = 5.3E-9). These results suggested that a robust humoral response and timely viral clearance (as seen in SubA compared to SubC), as well as better management of oxidative stress and anti-coagulation responses during hyperinflammation (SubE compared to SubF), may be crucial for improved post-acute recovery.

**Figure 6:**
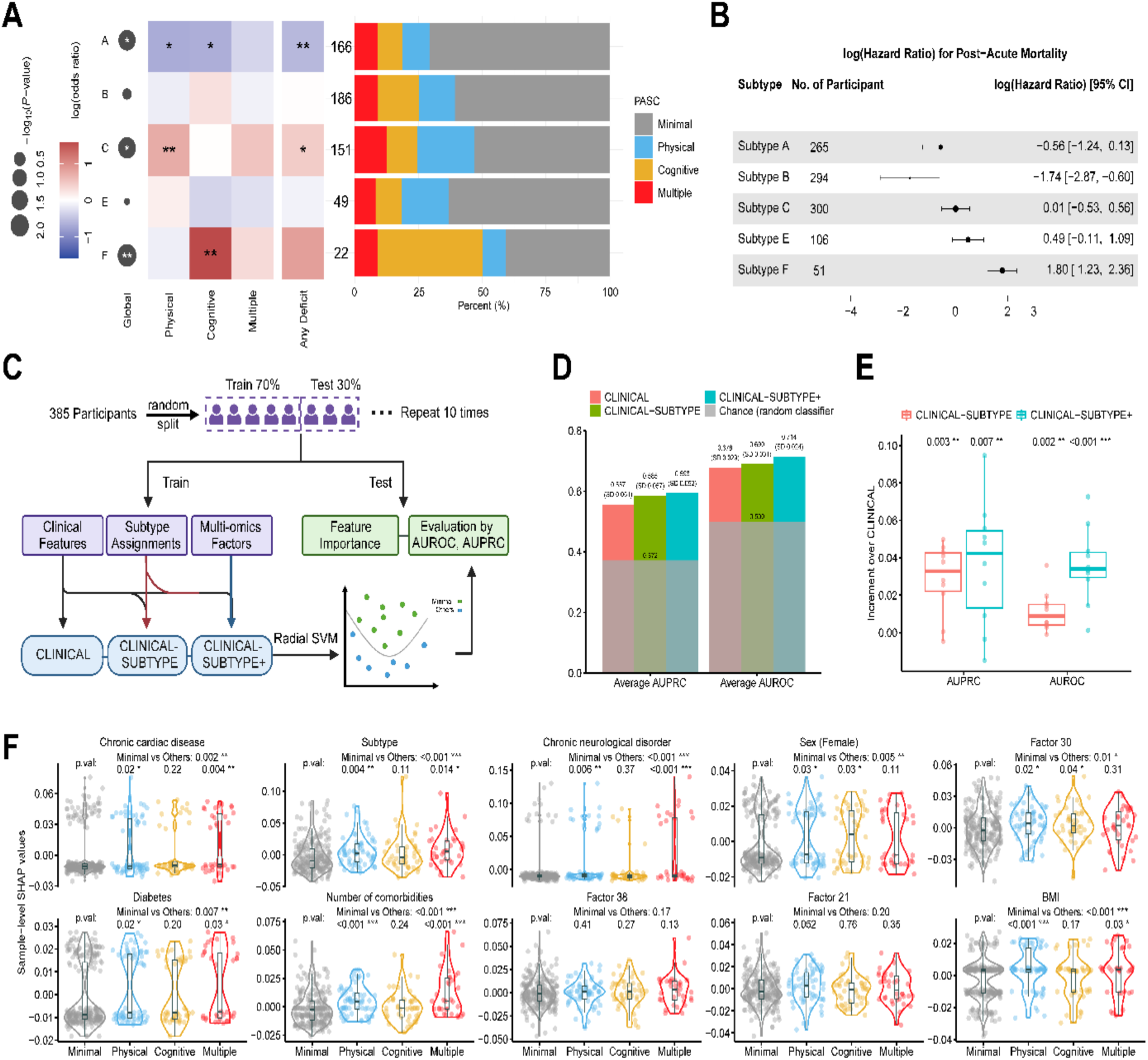
Machine learning models for PASC prediction. **(A)** Comparisons of PASC across subtypes using Fisher exact test. The overall p-value comparing all subtypes and all deficit groups was 6.4E-3. The leftmost row legend shows if the PASC groups were differentially enriched in a given subtype compared to other subtypes (Global p.val). Entries in the middle heatmap shows if each deficit PASC group (Physical, Cognitive, and Multiple) is enriched or not relative to minimal deficit in a given subtype, and the any deficit column examines the enrichment of all the deficit group relative to minimal group, with colors indicate log odds ratio relative to other group and significance annotated by asterisks. The rightmost barplot shows the empirical PASC group proportions within each subtype. **(B)** Forest plot showing the log-transformed hazard ratios for post-acute mortality for each subtype. Horizontal lines represent the 95% confidence intervals for hazard ratios from Cox proportional-hazards model controlling for age and sex. Intervals crossing the vertical line at 0 suggest non-significant differences compared to other subtypes. **(C)** The training and evaluation pipeline of PASC prediction where 30% of test data was excluded from the training or model tuning for a fair evaluation. **(D)** Average testing AUPRC and AUROC of CLINICAL, CLINICAL-SUBTYPE and CLINICAL-SUBTYPE+. **(E)** Increments in average testing AUPRC and AUROC of CLINICAL-SUBTYPE and CLINICAL-SUBTYPE+ over CLINICAL. P-values are based on the one-sided paired Wilcoxon Signed-Rank tests. **(F)** Sample-level SHAP values of top 10 features in CLINICAL-SUBTYPE+, colored by PASC groups. P-values are based on t-tests comparing each deficit group to the well-recovered Minimal group. (*p<=0.05, **p<=0.01, ***p<=0.001).

### Improving PASC Predictions Using the Molecular Subtypes

We developed machine learning models using radial kernel support vector machine (SVM) to predict PASC outcomes based on clinical characteristics and molecular profiles from the first-month post-admission. Our goal was to assess whether the subtype molecular profiles could improve the identification of PASC (cognitive, physical, and multiple deficit groups) versus minimal deficit among participants with survey data. Three prediction models were trained using different feature sets. The CLINICAL model used 22 features covering demographics (admission age, sex, BMI) and acute clinical characteristics, including length of stay, SOFA score, Spike IgG, N1-CT, the total number and presence of various comorbidities (hypertension, diabetes, chronic pulmonary disease, asthma, chronic cardiac disease, chronic kidney disease, malignant neoplasm, chronic neurological disorder, chronic liver disease, history of solid organ or bone marrow transplant, HIV, current or former smoking and/or vaping, drug or alcohol abuse or cannabis use, and number of other comorbidities), and the number of acute complications. For longitudinal observations such as Spike IgG, we applied an exponentially weighted moving average to all available visits of one participant with the most recent visit receiving the largest weight. We compared the CLINICAL model to the CLINICAL-SUBTYPE model which also included the probability of a participant being in each subtype to examine if our unbiased subtype assignment can enhance our prediction of PASC. Finally, we constructed the CLINICAL-SUBTYPE+ model that further incorporated the 40 multi-omics factors to fully exploit the molecular profiles.

We randomly split the 385 participants (62% minimal deficit and 38% any deficit) that had no missing clinical characteristics out of the 587 total participants with PASC assignments into a training set (70%) and a testing set (30%). We trained the three prediction models on the training set and evaluated them on the test set. We repeated this procedure 10 times and measured model performance using the average AUROC and average AUPRC on the 10 testing sets for improved stability (Figure 6C). Integrating the subtypes and multi-omics factors with the clinical measures enhanced the predictive accuracy, with CLINICAL, CLINICAL-SUBTYPE, and CLINICAL-SUBTYPE+ models achieving an average test AUROC of 0.678, 0.690, and 0.714, and an average test AUPRC of 0.557, 0.585, and 0.593, respectively (Figure 6D). The empirical p-values from a one-sided paired Wilcoxon rank test for improvement from CLINICAL-SUBTYPE and CLINICAL-SUBTYPE+ over the clinical model were 0.002 and 9.8e-4 for AUROC, and 0.003 and 0.007 for AUPRC (Figure 6E). These findings suggest that unbiased molecular subtypes and the derived multi-omics factors can provide valuable information for assessing post-acute stage outcomes beyond what can be determined using listed clinical characteristics. We identified the top ten features for the CLINICAL-SUBTYPE+ model using the SHAP values that measure a feature’s contribution to the prediction model^82^ (Figure 6F). Among them, chronic cardiac disease, subtype assignment scores, chronic neurological disorder, female sex, factor 30, diabetes, number of comorbidities, and BMI have significantly larger sample-level SHAP values when comparing the PASC group to the minimal deficit group. While most of the significance was due to the difference between minimal deficit and physical/multiple deficits, female sex and factor 30 have significantly larger SHAP values in physical/cognitive deficits than in minimal deficit (Figure 6F).

Given the significant elevation of Factor 30 in patients who later developed PASC (p.val = 4.49E-3 in early visits; p.val = 1.30E-3 in late visits, Extended Data 7A), we investigated the biological functions most closely associated with this factor. Factor 30 was primarily influenced by serum soluble proteins (COR = 0.85, Extended Data 7B), with only weak association from larger assays like PBMC and nasal transcriptomics (COR < 0.2). Key contributors to Factor 30 included elevated pro-inflammatory mediators, such as the Th-2 cytokines IL-13 and IL-33^83,84^, along with reduced IL-10RA as the strongest negative contributor (Extended Data 7B). The aggregated kinetic profile of top serum proteins was consistently elevated in PASC patients (Extended Data 7C). Increased Th-2 inflammation has been previously noted in PASC relative to controls at the post-acute stage, likely promoting tissue remodeling and immune dysregulation^10,85,86^. Despite weak influence, Factor 30 was enriched in several function classes across other assays (adj.p < 0.05, Extended Data 7D and Table S7), including elevated lipid-related pathways (n3 and n6 fatty acids, lysophospholipids, and long-chain monounsaturated fatty acids) and complement/coagulation proteins in proteomics, mirroring the changes observed in Factor 2 which was associated with SubC and SubF (Extended Data 7D-E). Factor 30 also revealed opposite trends in key inflammatory pathways between PBMC and nasal compartments (Extended Data 7D, 7F & 7G), with upregulated interferon responses in the nasal transcriptomics and a downregulated trend of these genes in PBMC. Interestingly, TNFA signaling via NFKB pathway was upregulated in the PBMC but no in the nasal, with leading-edge features included negative feedback inhibitory genes such as SOCS1, SOCS3, DUSP2, DUSP4, and SMAD3, possibly explaining the downregulation of downstream anti-viral and inflammatory signaling in PBMC^87–89^.

## Discussion

Our large-scale study of 1,148 hospitalized COVID-19 patients has revealed five distinct molecular subtypes with unique immune profiles, providing crucial insights into the interplay of immune heterogeneity after COVID-19 infection and recovery trajectories at both acute and post-acute stages. These findings represent an important advancement in our understanding of COVID-19 pathophysiology and highlight the potential for personalized treatment approaches based on molecular subtyping.

Among the severe subtypes (SubA-SubC), we observed important differences in both immune responses and clinical outcomes. SubC, in particular, displayed higher levels of organ disease markers, including EMT and myocardiopathy-related proteins, alongside increased tryptophan catabolism and acetylated peptides. These immune programs correlated with higher rates of cardiovascular and renal complications, even after controlling for factors such as age, sex, and comorbidities. In early visits, SubC also exhibited slower viral clearance and lower humoral responses, despite more pronounced T-cell activity, in contrast to SubA which displayed a controlled elevation of innate response, a stronger humoral response but lower T-cell activity. In the critical subtypes, SubF was marked by an early hyperinflammatory state and persistent immune imbalance, likely related to oxidative stress, including the simultaneous occurrence of hypercoagulation and seemingly silent anticoagulation regulation^90,91^, broad elevation of fatty acids and their derivatives with an increase in peroxidation products like 13-HODE and 9-HODE^69^ and in n6/n3 polyunsaturated fatty acid ratio^92^. These factors may exacerbate inflammation and contribute to SubF’s higher rate of anemia and chronic complications^93^.

Although using only acute molecular data, we found that SubC had worse post-acute outcomes compared to SubA-B, including a higher prevalence of PASC and a larger proportion of participants who failed to respond to post-hospitalization surveys which might also be related to poor post-acute recovery. Similarly, SubF exhibited a higher prevalence of PASC and post-acute mortality compared to SubE. Consistent with the prior study that reported reduced antibody titers, elevated nasal viral load, and serum FGF21 during acute hospitalization associated with PASC^32^, we observed reduced antibody response and slower clearance of viral loads in SubC relative to SubA-B, and elevated FGF21 (in SPmod3) in SubC and SubF relative to other subtypes of similar acute severity (Figure 3A, 3D, Extended Data 4E). Additionally, the overlap between biological functions found in PASC-associated subtypes in this study and those reported in the prior literature using convalescent immune profiles suggests that some dysregulations observed in PASC might begin during the acute phase. For example, complement and coagulation dysregulation has been observed in PASC patients for up to 6 months^94^, while we observed a highly imbalanced complement and coagulation state in SubF during the acute stage.

This large-scale study moves beyond marker identification for an individual clinical outcome and provides a holistic view of immune heterogeneity and clinical complications at the patient level, with our highlighted findings potentially advancing personalized treatments in COVID-19 infection. For instance, SubF’s hyperinflammatory and pro-coagulant profile may benefit from targeted anti-inflammatory and anticoagulation therapies early in the disease course, while SubC might require a different approach focused on boosting humoral immunity and addressing slow viral clearance. However, we acknowledge that the immediate translational application of our results is challenging given the extensive molecular assays required to define these subtypes. Nonetheless, the subtypes identified here can be leveraged to develop a smaller set of signatures from individual assays that can effectively capture the immune profiles of these subtypes. For example, we identified 75 serum protein signatures and 47 PBMC transcriptomic signatures for accurate subtype prediction, with AUROC scores of 0.913 and 0.807, respectively (Extended Data 8A). These predicted subtype probabilities showed consistent associations with PASC patterns (Extended Data 8B).

Despite the strengths of this study, there are several limitations: 1) Our cohort was limited to hospitalized, unvaccinated patients infected with the ancestral SARS-CoV-2 strain, potentially limiting generalizability to milder cases, vaccinated individuals, or infections with variants of concern. However, the investigation into the hospitalized cohort is of particular interest due to its diverse clinical complications and the associated insights into treatments of severe patients. 2) The self-reported PASC surveys contain a high rate of non-respondents, likely influenced by participants’ health during recovery. For example, a small percentage of participants dropped out due to recorded post-acute mortality, while other unobserved causes could account for the rest. This issue is prevalent in observational studies on long COVID or other infectious diseases with a long follow-up, where survey participation may strongly depend on the individual’s well-being. 3) Although we recorded the use of certain COVID-19 treatments such as remdesivir, monoclonal antibodies, and steroids, detailed information on treatment assignment and timing was not fully captured, limiting our ability to assess their causal impact on immune responses and clinical outcomes. This is also an issue for many other observational studies for severe infections. 4) While our study has combined a large number of longitudinal assays, there are questions in the observed immune profiles that remain unresolved, requiring additional targeted analysis and deeper investigations to fully understand them. For example, IL17A/C levels, normally associated with Type-3 responses and more neutrophils, were elevated in SubC (Figure 3D-E, Extended Data 4E), despite lower early CXCL8 and neutrophil frequencies relative to SubA-B.

Our analysis has also included attempts to alleviate limitations 2 and 3. We mitigated limitation 2 by investigating the non-response patterns in addition to PASC assignments (Extended Data 6C). We observed that SubC exhibited both higher PASC incidence and a greater proportion of survey non-respondents compared to SubA-B. Similarly, SubF had a greater rate of survey non-respondents and a higher prevalence of PASC than SubE. While the critical SubE appeared to have low PASC prevalence comparable to the severe SubA, it showed a high rate of non-respondents comparable to SubC. High rates of participant drop-out in observational long COVID studies can potentially lead to biased findings. The observed selective reduction in respondents (potentially due to illness) may help explain seemingly contradictory reports in the literature regarding acute severity and PASC prevalence^32,95^. By examining both PASC outcomes and participant drop-out to the convalescent surveys among different subtypes, our analysis offers robust insights into links from acute severity and immunophenotypes to long-term outcomes.

We have examined the impact of the presence of recorded medications during acute infection in our attempt to alleviate limitation 3. Remdesivir and antibiotics were the therapeutics most differentially used across patient subtypes. Particularly, remdesivir was most frequently used in SubA and least in SubC (Extended Data 8C, adj.p = 6.2E-9/6.9E-10), with elevated, though not significant, usage in SubE relative to SubF, implying a possible role in mitigating long COVID, which was also suggested by previous studies ^96,97^. Even after adjusting for age, sex, and remdesivir usage using multinomial logistic regression, the identified subtypes continued to show distinct post-recovery patterns consistent with the unadjusted results: SubA had the best recovery, measured by both reported PASC and missing survey rates, while SubC showed worst post-acute outcomes among the severe subtypes, and the critical subtype SubE better than the critical subtype SubF. This pattern persisted after adjusting for all differentially used medications (Extended Data 8D).

In conclusion, our identification of distinct molecular subtypes in hospitalized COVID-19 patients represents a significant step towards personalized medicine in infectious disease management. By linking acute immune responses to both acute and long-term outcomes, these findings not only enhance our understanding of COVID-19 pathophysiology but also provide clues for targeted interventions and improved patient care that can be employed during acute infection to prevent long COVID.

## Methods

### Participant enrollment and sample collection

This study explored data from 1,152 participants in the IMPACC observational cohort study^2,6,7,31^, which enrolled from 20 hospitals across 15 medical centers in the United States between May 5th, 2020 and March 19th, 2021. The end of this enrollment period represented the beginning of the emergence of the B.1.1.7 alpha variant, which was first recognized in the US in Dec 2020 and spread rapidly in Q1 of 2021; as such SARS-CoV-2 variants of concern were not widely represented in the cohort. Eligible participants were those hospitalized with SARS-CoV-2 infection confirmed by reverse transcription polymerase chain reaction (RT-PCR) and symptoms or signs consistent with COVID-19. The detailed study design and schedule for clinical data and biologic sample collection, and shared core platform assessments were previously described^6,7^. Detailed clinical assessments and sampling of blood and upper respiratory tract were performed within approximately 72 hours of hospitalization (Visit 1), and on approximately Days 4 (Visit 2), 7 (Visit 3), 14 (Visit 4), 21 (Visit 5), 28 (Visit 6) after hospital admission (which corresponded to Visits 2 to 6), amounting 3,077 sampling events. As previously described^31^, biological sample collection and processing followed a standard protocol utilized by every participating academic institution. More details regarding the study cohort, sample processing and batch correction can be found in Supplementary Materials.

### Ethics statement

NIAID staff conferred with the Department of Health and Human Services Office for Human Research Protections (OHRP) regarding potential applicability of the public health surveillance exception [45CFR46.102] to the IMPACC study protocol. OHRP concurred that the study satisfied criteria for the public health surveillance exception, and the IMPACC study team sent the study protocol, and participant information sheet for review, and assessment to institutional review boards (IRBs) at participating institutions. Twelve institutions elected to conduct the study as public health surveillance, while 3 sites with prior IRB-approved biobanking protocols elected to integrate and conduct IMPACC under their institutional protocols (University of Texas at Austin, IRB 2020-04-0117; University of California San Francisco, IRB 20-30497; Case Western Reserve University, IRB STUDY20200573) with informed consent requirements. Participants enrolled under the public health surveillance exclusion were provided information sheets describing the study, samples to be collected, and plans for data de-identification, and use. Those that requested not to participate after reviewing the information sheet were not enrolled. In addition, participants did not receive compensation for study participation while inpatient, and subsequently were offered compensation during outpatient follow-ups.

### Study design rationale and overview

A subset of the IMPACC cohort has previously been analyzed to identify immune correlates or multi-omics immune programs predictive and associated with acute mortality, respiratory severity, and specific demographic or clinical characteristics such as aging^6,7,98^. However, while its large data volume enabled us to reach more conclusive statements identifying immune profiles associated with specific clinical endpoints of interest, this rich and large-scale database also offers an underexplored opportunity to conduct deep analysis on immune heterogeneity and identify immune-based patient subtypes. These subtypes can potentially guide us to gain more comprehensive insights into the host-responses and allow us to understand their links to a wide range of clinical outcomes, both from the acute stage and the convalescent phase, in a unified framework.

Following this goal, this study processed datasets encompassing plasma proteomics (global and targeted; PPG and PPT, respectively), serum proteins (SPT), plasma metabolites (PMG), and mRNAs from nasal swabs and PBMCs (NGX and PGX, respectively) were utilized to construct multi-omics factors. These factors were generated using the Multiple Co-Inertia Analysis (MCIA) model to capture major covariation patterns across omics datasets. Patient-level subtypes were then defined through hierarchical clustering based on averaged pair-wise distances across all available visits.

The associations of these subtypes with a broad spectrum of demographic and clinical outcomes—including acute mortality, respiratory severity, non-respiratory complications, post-acute mortality, and long COVID—were systematically examined. Additionally, multi-omics factors were integrated with measurements of nasal viral load, serum SARS-CoV-2 antibody titers, and whole blood CyTOF data from the same cohort to investigate immune programs distinguishing the identified subtypes.

Further details regarding multi-omics construction, subtype identification, and the investigation of demographic and clinical outcomes, as well as the immunological characterization of the subtypes, are provided in the Supplementary Materials.

### Statistics

#### Multi-omics annotation

High-contributing features for each assay were defined as the features whose absolute coefficients (from regressing features to factors) >=0.3 and adjusted p-values of significance p.adj <= 0.01 to identify the top contributing features associated with MCIA multi-omics factors. Functional enrichment analysis for multi-omics factors was performed using high-contributing features with minimum hypergeometric test (mHG) and a variety of publicly available databases, with only those adj.p < 0.05 considered. The pathway activities were calculated as a weighted sum of a selected set of features in a pathway.

#### Hypothesis tests

Multiple comparisons were accounted for via Benjamini-Hochberg correction by default, with adjusted p-values (adj.p) < 0.05 considered significant unless otherwise stated. Generalized linear regression was utilized when assessing the association between subtypes and demographic or clinical measures, where linear regression was used for baseline neutrophil and lymphocyte counts, ordinal regressions was used for ordinal TG group and age quantile group, cox proportional hazard model was used for mortality evaluation, and logistic regression was used for other categorical measures. Except for evaluating the demographic association, basic demographics (age and sex) have been adjusted in other hypothesis testing analysis. Additional examination of the association between subtypes and clinical complications after adjusting for significant comorbidities has been conducted to account for the influence of prior medical conditions on complications. Generalized mixed-effect modeling was utilized to investigate differential kinetics of immune profiles (including multi-omics factors, cytokine modules, pathway activities, viral load, SARS-CoV-2 specific IgG and cell frequencies measured by CyTOF) across subtypes using samples from all visits after further adjusting for participant id as random-effect. Furthermore, a linear mixed effect model was applied to obtain the differential patterns with directionality comparing subtypes in early visits (visit 1-visit 3) and late visits (visit 3-visit 6), after adjusting for admission date, sex, age as fixed effects and participant id and enrollment site as the random effects unless otherwise specified. Such a linear mixed effect modeling is also utilized to investigate the relationship between viral load and inflammatory multi-omics factors/serum soluble protein modules.

#### Machine learning model construction for PASC prediction and feature importance rank

For longitudinal features which samples from multiple visits for a participant (including N1-CT, Spike IgG, and the 40 multi-omics factors), we applied an exponentially weighted moving average to all available visits of one participant, in which the most recent visit was given the largest weight. We also constructed the subtype assignment scores based on the Euclidean distance from each participant to the subtype centers. After the feature engineering, we fitted a radial kernel SVM on the training set. We applied a 5-fold cross-validation using the AUROC as the metric to determine the tuning parameters. Then, we used the SHAP value^82^ to evaluate the importance of each feature in the CLNICAL-SUBTYPE+ model. For each of the top 10 features, we did a two-sample one-sided t-test on the sample-level SHAP values (evaluated on the test samples) for minimal deficit (set to be the lower level) versus physical deficit, cognitive deficit, multiple deficits, and any deficit, respectively.

#### Sparse models for subtype prediction

We developed models for subtype prediction using selected sparse markers from targeted proteomics, global proteomics, serum olink, plasma metabolomics, nasal transcriptomics, and PBMC transcriptomics. Specifically, we selected markers by generalized LASSO linear regression model and then fitted a radial SVM using the selected ones. We then tested the associations between the predicted subtype probabilities and PASC patterns using multinomial and binomial regressions.

See Supplementary Materials for detailed descriptions of all statistical analyses and models.

## Author contributions

Conceptualization: IMPACC Network, EM, HK, HS, BP, CBC, ADA, PMB, SF, CP, BP, SHK, OL, MCA, AI, JDA, LIRE, LG

Formal analysis: KW, YN, CM, CS, HS, MK, JP, LL, AH, NDJ, LG

Software: KW, YN, CM, LG Methodology: LG

Resources: SB, SKS, FK, LR, HvB, MW, HS, WE, CL, OL, MCA, HM, RRM

Funding Acquisition: IMPACC Network Supervision: OL, MCA, AI, JDA, LIRE, LG

All authors wrote, edited, and reviewed the manuscript.

## Data availability

Data used in this study is available at ImmPort Shared Data under the accession number SDY1760 (https://www.immport.org/shared/search?text=SDY1760%20) and in the NLM’s Database of Genotypes and Phenotypes (dbGaP) under the accession number phs002686.v2.p2 (https://www.ncbi.nlm.nih.gov/projects/gap/cgi-bin/study.cgi?study_id=phs002686.v2.p2).

## Code availability

All code is deposited on Bitbucket repository (https://bitbucket.org/kleinstein/impacc-public-code/src/master/molecular_subtyping).

## #The IMPACC Network

**National Institute of Allergy and Infectious Diseases, National Institute of Health, Bethesda, MD 20814, USA:** Patrice M. Becker, Alison D. Augustine, Steven M. Holland, Lindsey B. Rosen, Serena Lee, Tatyana Vaysman

**Clinical and Data Coordinating Center (CDCC) Precision Vaccines Program, Boston Children’s Hospital, Boston, MA 02115, USA:** Al Ozonoff, Joann Diray-Arce, Jing Chen, Alvin Kho, Carly E. Milliren, Annmarie Hoch, Ana C. Chang, Kerry McEnaney, Brenda Barton, Claudia Lentucci, Maimouna D. Murphy, Mehmet Saluvan, Tanzia Shaheen, Shanshan Liu, Caitlin Syphurs, Marisa Albert, Arash Nemati Hayati, Robert Bryant, James Abraham, Sanya Thomas, Mitchell Cooney, Meagan Karoly

**Benaroya Research Institute, University of Washington, Seattle, WA 98101, USA:** Matthew C. Altman, Naresh Doni Jayavelu, Scott Presnell, Bernard Kohr, Tomasz Jancsyk, Azlann Arnett

**La Jolla Institute for Immunology, La Jolla, CA 92037, USA:** Bjoern Peters, James A. Overton, Randi Vita, Kerstin Westendorf

**Knocean Inc. Toronto, ON M6P 2T3, Canada:** James A. Overton

**Precision Vaccines Program, Boston Children’s Hospital, Harvard Medical School, Boston, MA 02115, USA:** Ofer Levy, Hanno Steen, Patrick van Zalm, Benoit Fatou, Kinga K. Smolen, Arthur Viode, Simon van Haren, Meenakshi Jha, David Stevenson, Oludare Odumade

**Brigham and Women’s Hospital, Harvard Medical School, Boston, MA 02115, USA:** Lindsey R. Baden, Kevin Mendez, Jessica Lasky-Su, Alexandra Tong, Rebecca Rooks, Michael Desjardins, Amy C. Sherman, Stephen R. Walsh, Xhoi Mitre, Jessica Cauley, Xiofang Li, Bethany Evans, Christina Montesano, Jose Humberto Licona, Jonathan Krauss, Nicholas C. Issa, Jun Bai Park Chang, Natalie Izaguirre

**Metabolon Inc, Morrisville, NC 27560, USA:** Scott R. Hutton, Greg Michelotti, Kari Wong

**Prevention of Organ Failure (PROOF) Centre of Excellence, University of British Columbia, Vancouver, BC V6T 1Z3, Canada:** Scott J. Tebbutt, Casey P. Shannon

**Case Western Reserve University and University Hospitals of Cleveland, Cleveland, OH 44106, USA:** Rafick-Pierre Sekaly, Slim Fourati, Grace A. McComsey, Paul Harris, Scott Sieg, Susan Pereira Ribeiro

**Drexel University, Tower Health Hospital, Philadelphia, PA 19104, USA:** Charles B. Cairns, Elias K. Haddad, Michele A. Kutzler, Mariana Bernui, Gina Cusimano, Jennifer Connors, Kyra Woloszczuk, David Joyner, Carolyn Edwards, Edward Lee, Edward Lin, Nataliya Melnyk, Debra L. Powell, James N. Kim, I. Michael Goonewardene, Brent Simmons, Cecilia M. Smith, Mark Martens, Brett Croen, Nicholas C. Semenza, Mathew R. Bell, Sara Furukawa, Renee McLin, George P. Tegos, Brandon Rogowski, Nathan Mege, Kristen Ulring, Pam Schearer, Judie Sheidy, Crystal Nagle

**MyOwnMed Inc., Bethesda, MD 20817, USA:** Vicki Seyfert-Margolis

**Emory School of Medicine, Atlanta, GA 30322, USA:** Nadine Rouphael, Steven E. Bosinger, Arun K. Boddapati, Greg K. Tharp, Kathryn L. Pellegrini, Brandi Johnson, Bernadine Panganiban, Christopher Huerta, Evan J. Anderson, Hady Samaha, Jonathan E. Sevransky, Laurel Bristow, Elizabeth Beagle, David Cowan, Sydney Hamilton, Thomas Hodder, Amer Bechnak, Andrew Cheng, Aneesh Mehta, Caroline R. Ciric, Christine Spainhour, Erin Carter, Erin M. Scherer, Jacob Usher, Kieffer Hellmeister, Laila Hussaini, Lauren Hewitt, Nina Mcnair, Susan Pereira Ribeiro, Sonia Wimalasena

**Icahn School of Medicine at Mount Sinai, New York, NY 10029, USA:** Ana Fernandez-Sesma, Viviana Simon, Florian Krammer, Harm Van Bakel, Seunghee Kim-Schulze, Ana Silvia Gonzalez Reiche, Jingjing Qi, Brian Lee, Juan Manuel Carreño, Gagandeep Singh, Ariel Raskin, Johnstone Tcheou, Zain Khalil, Adriana van de Guchte, Keith Farrugia, Zenab Khan, Geoffrey Kelly, Komal Srivastava, Lily Q. Eaker, Maria C. Bermúdez-González, Lubbertus C.F. Mulder, Katherine F. Beach, Miti Saksena, Deena Altman, Erna Kojic, Levy A. Sominsky, Arman Azad, Dominika Bielak, Hisaaki Kawabata, Temima Yellin, Miriam Fried, Leeba Sullivan, Sara Morris, Giulio Kleiner, Daniel Stadlbauer, Jayeeta Dutta, Hui Xie, Manishkumar Patel, Kai Nie

**Immunai Inc. New York, NY 10016, USA:** Adeeb Rahman

**Oregon Health Sciences University, Portland, OR 97239, USA:** William B. Messer, Catherine L. Hough, Sarah A.R. Siegel, Peter E. Sullivan, Zhengchun Lu, Amanda E. Brunton, Matthew Strand, Zoe L. Lyski, Felicity J. Coulter, Courtney Micheleti

**Stanford University School of Medicine, Palo Alto, CA 94305, USA:** Holden Maecker, Bali Pulendran, Kari C. Nadeau, Yael Rosenberg-Hasson, Michael Leipold, Natalia Sigal, Angela Rogers, Andrea Fernandes, Monali Manohar, Evan Do, Iris Chang, Alexandra S. Lee, Catherine Blish, Henna Naz Din, Jonasel Roque, Linda N. Geng, Maja Artandi, Mark M. Davis, Neera Ahuja, Samuel S. Yang, Sharon Chinthrajah, Thomas Hagan

**David Geffen School of Medicine at the University of California Los Angeles, Los Angeles CA 90095, USA:** Elaine F. Reed, Joanna Schaenman, Ramin Salehi-Rad, Adreanne M. Rivera, Harry C. Pickering, Subha Sen, David Elashoff, Dawn C. Ward, Jenny Brook, Estefania Ramires Sanchez, Megan Llamas, Claudia Perdomo, Clara E. Magyar, Jennifer Fulcher

**University of California San Francisco, San Francisco, CA 94115, USA:** David J. Erle, Carolyn S. Calfee, Carolyn M. Hendrickson, Kirsten N. Kangelaris, Viet Nguyen, Deanna Lee, Suzanna Chak, Rajani Ghale, Ana Gonzalez, Alejandra Jauregui, Carolyn Leroux, Luz Torres Altamirano, Ahmad Sadeed Rashid, Andrew Willmore, Prescott G. Woodruff, Matthew F. Krummel, Sidney Carrillo, Alyssa Ward, Charles R. Langelier, Ravi Patel, Michael Wilson, Ravi Dandekar, Bonny Alvarenga, Jayant Rajan, Walter Eckalbar, Andrew W. Schroeder, Gabriela K. Fragiadakis, Alexandra Tsitsiklis, Eran Mick, Yanedth Sanchez Guerrero, Christina Love, Lenka Maliskova, Michael Adkisson, Aleksandra Leligdowicz, Alexander Beagle, Arjun Rao, Austin Sigman, Bushra Samad, Cindy Curiel, Cole Shaw, Gayelan Tietje-Ulrich, Jeff Milush, Jonathan Singer, Joshua J. Vasquez, Kevin Tang, Legna Betancourt, Lekshmi Santhosh, Logan Pierce, Maria Tecero Paz, Michael Matthay, Neeta Thakur, Nicklaus Rodriguez, Nicole Sutter, Norman Jones, Pratik Sinha, Priya Prasad, Raphael Lota, Saurabh Asthana, Sharvari Bhide, Tasha Lea, Yumiko Abe-Jones

**Yale School of Medicine, New Haven, CT 06510, USA:** David A. Hafler, Ruth R. Montgomery, Albert C. Shaw, Steven H. Kleinstein, Jeremy P. Gygi, Shrikant Pawar, Anna Konstorum, Ernie Chen, Chris Cotsapas, Xiaomei Wang, Leqi Xu, Charles Dela Cruz, Akiko Iwasaki, Subhasis Mohanty, Allison Nelson, Yujiao Zhao, Shelli Farhadian, Hiromitsu Asashima, Omkar Chaudhary, Andreas Coppi, John Fournier, M. Catherine Muenker, Allison Nelson, Khadir Raddassi, Michael Rainone, William Ruff, Syim Salahuddin, Wade L. Shulz, Pavithra Vijayakumar, Haowei Wang, Esio Wunder Jr., H. Patrick Young, Albert I. Ko, Xiomei Wang

**Yale School of Public Health, New Haven, CT 06510, USA:** Denise Esserman, Leying Guan, Anderson Brito, Jessica Rothman, Nathan D. Grubaugh

**Baylor College of Medicine and the Center for Translational Research on Inflammatory Diseases, Houston, TX 77030, USA:** David B. Corry, Farrah Kheradmand, Li-Zhen Song, Ebony Nelson

**Oklahoma University Health Sciences Center, Oklahoma City, OK 73104, USA**: Jordan P. Metcalf, Nelson I. Agudelo Higuita, Lauren A. Sinko, J. Leland Booth, Douglas A. Drevets, Brent R. Brown

**University of Arizona, Tucson AZ 85721, USA:** Monica Kraft, Chris Bime, Jarrod Mosier, Heidi Erickson, Ron Schunk, Hiroki Kimura, Michelle Conway, Dave Francisco, Allyson Molzahn, Connie Cathleen Wilson, Ron Schunk, Trina Hughes, Bianca Sierra

**University of Florida, Gainesville, FL 32611, USA:** Mark A. Atkinson, Scott C. Brakenridge, Ricardo F. Ungaro, Brittany Roth Manning, Lyle Moldawer University of Florida, Jacksonville, FL 32218, USA: Jordan Oberhaus, Faheem W. Guirgis University of South Florida, Tampa FL 33620, USA: Brittney Borresen, Matthew L. Anderson

**University of Texas, Austin, TX 78712, USA:** Lauren I. R. Ehrlich, Esther Melamed, Cole Maguire, Dennis Wylie, Justin F. Rousseau, Kerin C. Hurley, Janelle N. Geltman, Nadia Siles, Jacob E. Rogers.

## IMPACC Network Competing Interests

The Icahn School of Medicine at Mount Sinai has filed patent applications relating to SARS-CoV-2 serological assays, NDV-based SARS-CoV-2 vaccines influenza virus vaccines and influenza virus therapeutics which list Florian Krammer as co-inventor. Mount Sinai has spun out a company, Kantaro, to market serological tests for SARS-CoV-2 and another company, Castlevax, to develop SARS-CoV-2 vaccines. Florian Krammer is co-founder and scientific advisory board member of Castlevax. Florian Krammer has consulted for Merck, Curevac, Seqirus, GSK and Pfizer and is currently consulting for 3rd Rock Ventures, Sanofi, Gritstone and Avimex. The Krammer laboratory is also collaborating with Dynavax on influenza vaccine development and with VIR on influenza virus therapeutics development. Viviana Simon is a co-inventor on a patent filed relating to SARS-CoV-2 serological assays (the “Serology Assays”). Ofer Levy is a named inventor on patents held by Boston Children’s Hospital relating to vaccine adjuvants and human in vitro platforms that model vaccine action. His laboratory has received research support from GlaxoSmithKline (GSK) and is a co-founder of and advisor to Ovax, Inc. Charles Cairns serves as a consultant to bioMerieux and is funded for a grant from Bill & Melinda Gates Foundation. James A Overton is a consultant at Knocean Inc. Jessica Lasky-Su serves as a scientific advisor of Precion Inc. Scott R. Hutton, Greg Michelloti and Kari Wong are employees of Metabolon Inc. Vicki Seyfer-Margolis is a current employee of MyOwnMed. Nadine Rouphael reports grants or contracts with Merck, Sanofi, Pfizer, Vaccine Company and Immorna, and has participated on data safety monitoring boards for Moderna, Sanofi, Seqirus, Pfizer, EMMES, ICON, BARDA, and CyanVan, Imunon Micron. N.R. has also received support for meetings/travel from Sanofi and Moderna and honoraria from Virology Education and Krog consulting. Adeeb Rahman is a current employee of Immunai Inc. Steven Kleinstein is a consultant related to ImmPort data repository for Peraton. Nathan Grabaugh is a consultant for Tempus Labs and the National Basketball Association. Akiko Iwasaki is a consultant for 4BIO, Blue Willow Biologics, Revelar Biotherapeutics, RIGImmune, Xanadu Bio, Paratus Sciences. Monika Kraft receives research funds paid to her institution from NIH, ALA; Sanofi, Astra-Zeneca for work in asthma, serves as a consultant for Astra-Zeneca, Sanofi, Chiesi, GSK for severe asthma; is a co-founder and CMO for RaeSedo, Inc, a company created to develop peptidomimetics for treatment of inflammatory lung disease. Esther Melamed received research funding from Babson Diagnostics and honorarium from Multiple Sclerosis Association of America and has served on the advisory boards of Genentech, Horizon, Teva, and Viela Bio. Carolyn Calfee receives research funding from NIH, FDA, DOD, Roche-Genentech and Quantum Leap Healthcare Collaborative as well as consulting services for Janssen, Vasomune, Gen1e Life Sciences, NGMBio, and Cellenkos. Wade Schulz was an investigator for a research agreement, through Yale University, from the Shenzhen Center for Health Information for work to advance intelligent disease prevention and health promotion; collaborates with the National Center for Cardiovascular Diseases in Beijing; is a technical consultant to Hugo Health, a personal health information platform; cofounder of Refactor Health, an AI-augmented data management platform for health care; and has received grants from Merck and Regeneron Pharmaceutical for research related to COVID-19. Grace A McComsey received research grants from Rehdhill, Cognivue, Pfizer, and Genentech, and served as a research consultant for Gilead, Merck, Viiv/GSK, and Janssen. Linda N. Geng received research funding paid to her institution from Pfizer, Inc.

## Funding

NIH (3U01AI167892-03S2, 3U01AI167892-01S2, 5R01AI135803-03, 5U19AI118608-04, 5U19AI128910-04, 4U19AI090023-11, 4U19AI118610-06, R01AI145835-01A1S1, 5U19AI062629-17, 5U19AI057229-17, 5U19AI057229-18, 5U19AI125357-05, 5U19AI128913-03, 3U19AI077439-13, 5U54AI142766-03, 5R01AI104870-07, 3U19AI089992-09, 3U19AI128913-03, and 5T32DA018926-18); NIAID, NIH (3U19AI1289130, U19AI128913-04S1, and R01AI122220); NCATS, NIH UM1TR004528 and National Science Foundation (DMS2310836).

## Acknowledgment

We thank the participants of the study for their voluntary enrollment and contribution of samples for this work. See the supplement for details on the IMPACC Network. We acknowledge the assistance of the following individuals: Sanya Thomas, Mitchell Cooney, Shun Rao, Sofia Vignolo, and Elena Morrocchi (all from the CDCC); Arash Naeim, Marianne Bernardo, Sarahmay Sanchez, Shannon Intluxay, Clara Magyar, Jenny Brook, Estefania Ramires-Sanchez, Megan Llamas, Claudia Perdomo, Clara E. Magyar, and Jennifer A. Fulcher (all from the David Geffen School of Medicine at UCLA); members of the UCLA Center for Pathology Research Services and the Pathology Research Portal; M. Catherine Muenker, Dimitri Duvilaire, Maxine Kuang, William Ruff, Khadir Raddassi, Denise Shepherd, Haowei Wang, Omkar Chaudhary, Syim Salahuddin, John Fournier, Michael Rainone, and Maxine Kuang (all from the Yale School of Medicine). We thank the leadership of Boston Children’s Hospital including Drs. Wendy Chung, Gary Fleisher and Kevin Churchwell for their support for the Precision Vaccines Program. Dr. Augustine’s and Becker’s co-authorship of this report does not necessarily represent the official views of the National Institute of Allergy and Infectious Diseases, the National Institutes of Health or any other agency of the United States Government.

---

**Extended Data Figure 1:**
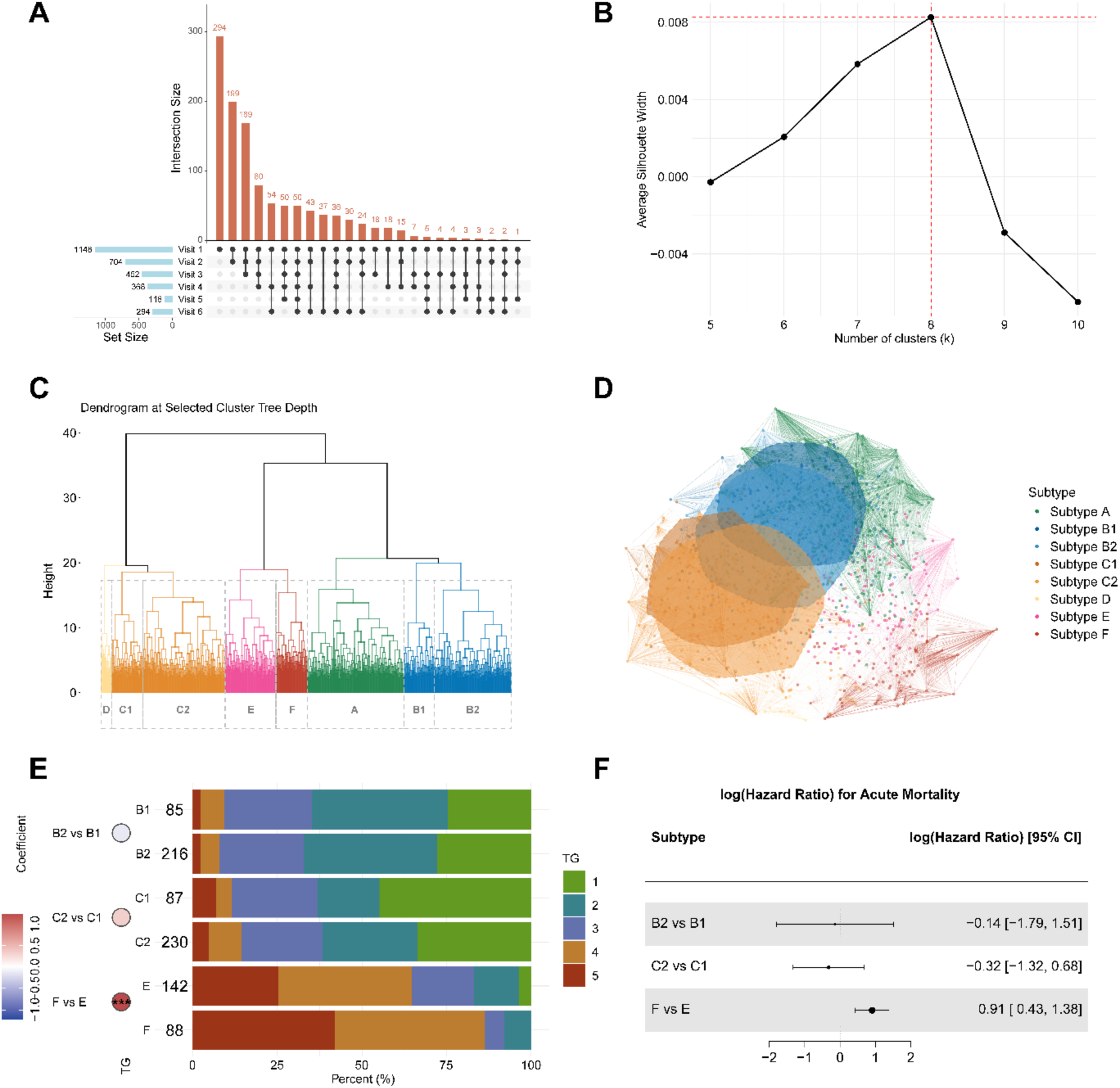
Participant availability across visits and subtype merging following hierarchical clustering. **(A)** Upset plot depicting the number of participants from visit 1 through visit 6 utilized for subtype construction (n=1,148). **(B)** Line chart illustrating the average silhouette width for selecting the number of clusters ranging from 5 to 10 in hierarchical clustering. The optimal number of clusters directly determined from hierarchical clustering is 8 where the silhouette width is maximized. **(C)** Cluster dendrogram explaining the hierarchical relationship among 8 clusters. **(D)** Network plot illustrating the multi-dimensional scaling of the distance between participants. Each ellipse encompasses 60% of the points from the centroid of its respective cluster, which revealed that the two most adjacent leaf clusters B1 and B2 are highly overlapping (in dark and light yellow) and the two most adjacent leaf clusters C1 and C2 are highly overlapping (in dark and light blue). **(E)** Barplot colored by TG showing comparisons between three pairs of closest adjacent leaf clusters (B1, B2), (C1, C2), and (E, F) through ordinal regression test after adjusting for admission age and sex. Significance was calculated via the ordinal regression model. (*p<=0.05, **p<=0.01, ***p<=0.001). **(F)** Forest plot showing the log-transformed hazard ratios for acute mortality comparing within three pairs of adjacent leaf clusters (B1, B2), (C1, C2), (E, F). Horizontal lines represent the 95% confidence intervals for log hazard ratios from Cox proportional-hazards model controlling for age and sex. Intervals crossing the vertical line at 0 suggest non-significant differences between the two groups.

**Extended Data Figure 2:**
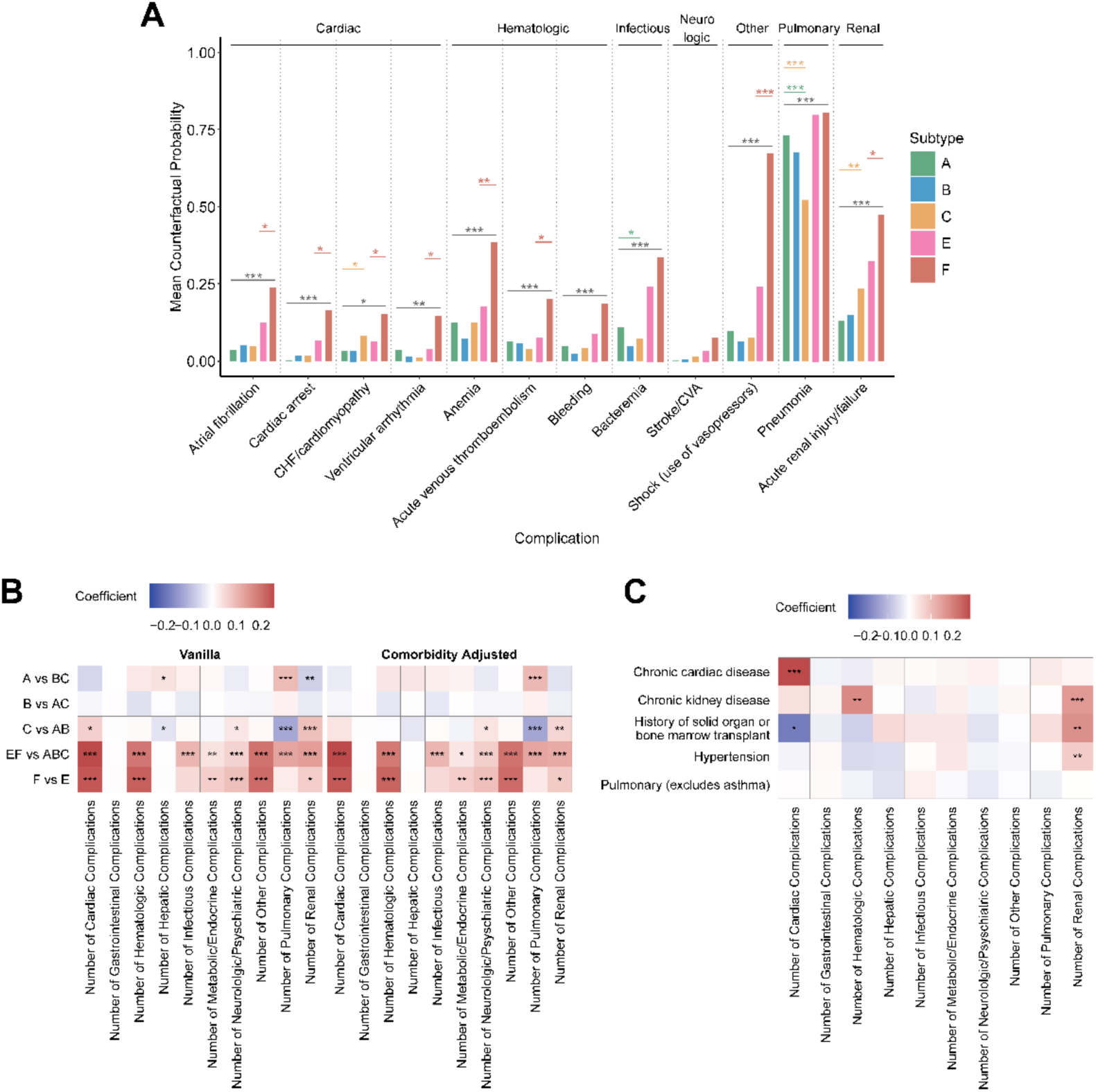
Molecular subtypes exhibit distinct complications grouped by bodily systems and functions. **(A)** Barplot showing mean counterfactual probability of significant complications (Chi-squared test adj.p<=0.001) characterized by bodily systems and functions. The mean probability isolates the contribution of subtypes from the logistic regression, which models group-wise comparison of complications after controlling for age, sex, and 5 significant comorbidities. Horizontal lines marked with asterisks show the significance after multiple test correction, with the grey line indicating the comparison between EF and ABC. Lines and asterisks matching the colors of A, B, and C denote comparisons of each group against the other two – A vs BC, B vs AC, and C vs AB respectively. The red line matching the color of F represents the comparison between F and E. **(B)** Heatmap of complication categories for group-wise subtype comparison using linear regression, showing both the version adjusted for age and sex and the version further adjusted for significant comorbidities. **(C)** Heatmap of complication categories against 5 significant comorbidities. Coefficients and significance were derived from the same regression model used in the comorbidity-adjusted analysis in (B). (*adj.p<=0.05, **adj.p<=0.01, ***adj.p<=0.001).

**Extended Data Figure 3:**
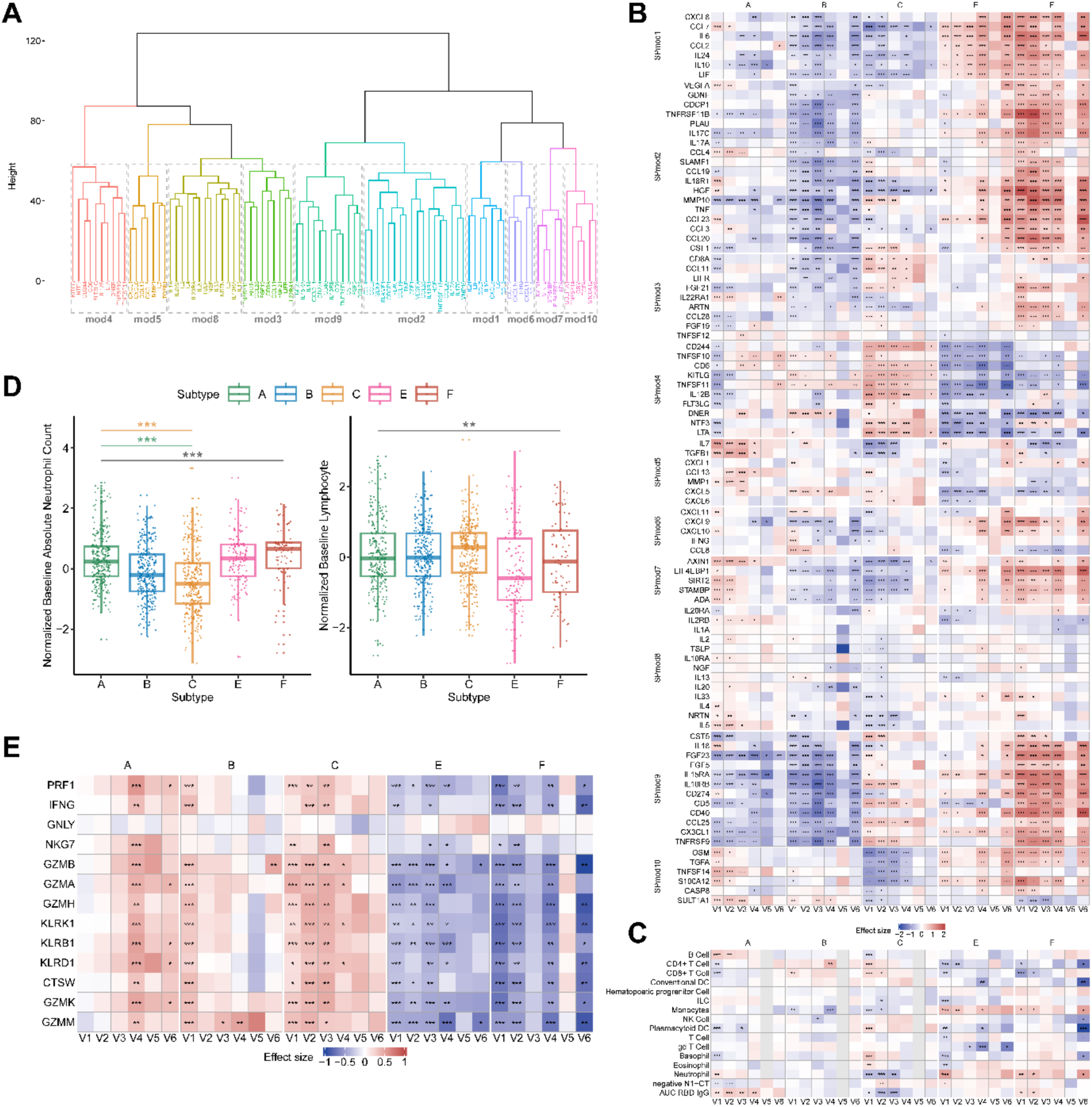
**(A)** Hierarchical clustering dendrogram for grouping serum soluble protein profiles into 10 modules. **(B)** Heatmap comparing the serum protein profiles of one subtype against others using t-test. **(C)** Heatmap comparing one subtype vs others in cell composition, viral load and antibody using t-test. **(D)** Barplot of baseline absolute neutrophil count and lymphocyte, with the grey line indicating the comparison between EF and ABC. Lines and asterisks matching the colors of A, B, and C denote comparisons of each group against the other two – A vs BC, B vs AC, and C vs AB respectively. **(E)** Heatmap comparing T cell cytotoxic functions of one subtype against others during visits 1-3 and visits 4-6 using t-test. (*adj.p<=0.05, **adj.p<=0.01, ***adj.p<=0.001).

**Extended Data Figure 4:**
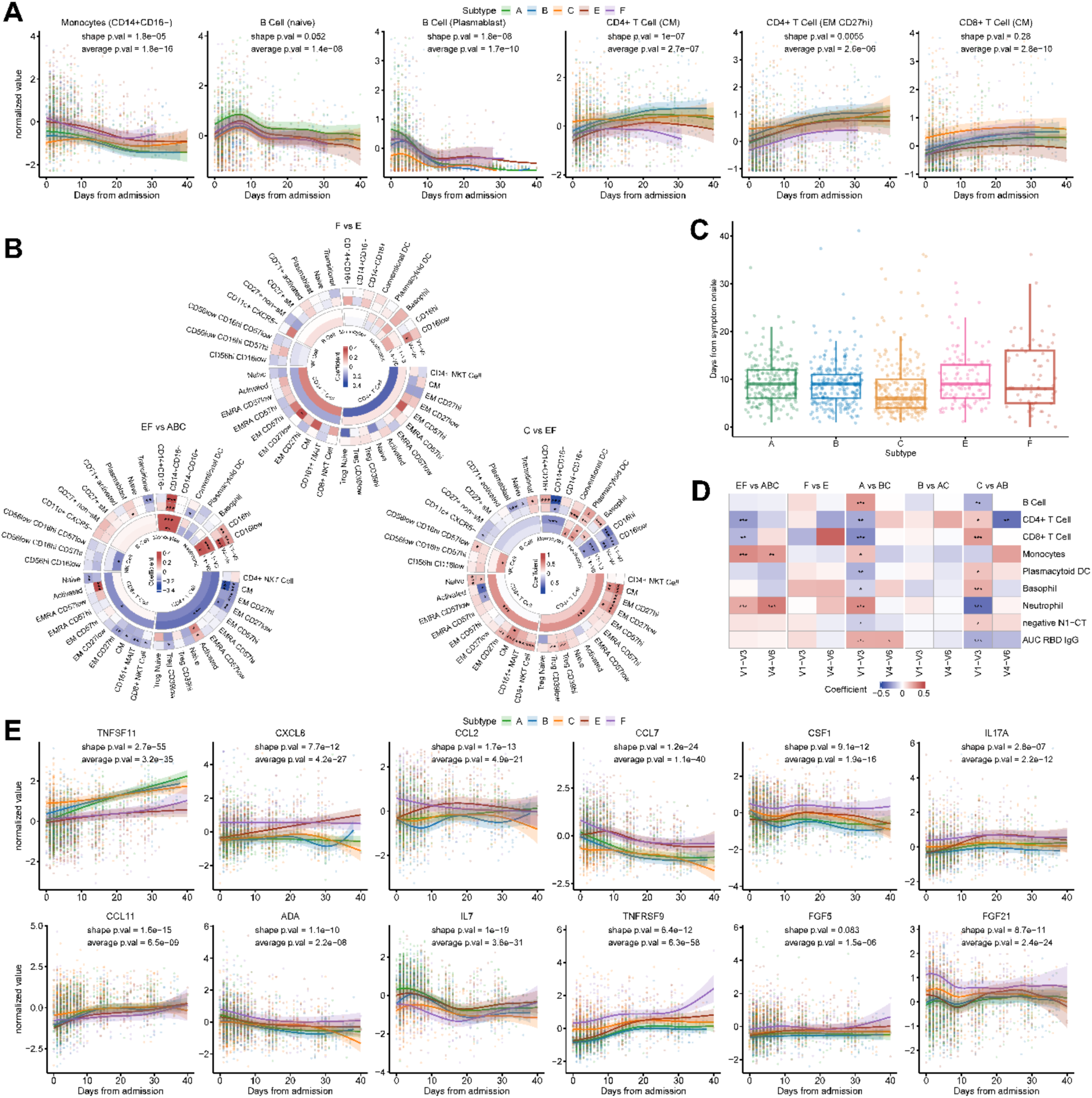
**(A)** Longitudinal trajectories of CD14+CD16-monocytes, naive B cell, plasmablast B cell, CM CD4+ T cell, EM CD27hi CD4+ T cell and CM CD8+ T cell, colored by subtypes. Shaded region denotes 95% confidence interval from generalized additive mixed model of the fitted trajectory (thick line). **(B)** Circular heatmaps of comparisons in whole blood (CyTOF) of both parent and child populations for visits 1-3 and visits 4-6 between SubEF and SubABC, SubF and SubE, SubC and SubEF. **(C)** Boxplot of days from symptom onsite for major subtypes. **(D)** Heatmap of comparisons in cell composition, viral load and antibody for visits 1-3 and visits 4-6 among 5 major subtypes using linear mixed effect modeling after further adjusting for participant ID and enrollment site as random effects and age, sex, and days from symptom onset as fixed effects. **(E)** Longitudinal trajectories of TNFSF11, CXCL8, CCL2, CCL7, CSF1, IL17A, CCL11, ADA, IL7, TNFRSF9, FGF5, and FGF21, colored by subtypes. Shaded region denotes 95% confidence interval from generalized additive mixed model of the fitted trajectory (thick line). (*adj.p<=0.05, **adj.p<=0.01, ***adj.p<=0.001).

**Extended Data Figure 5:**
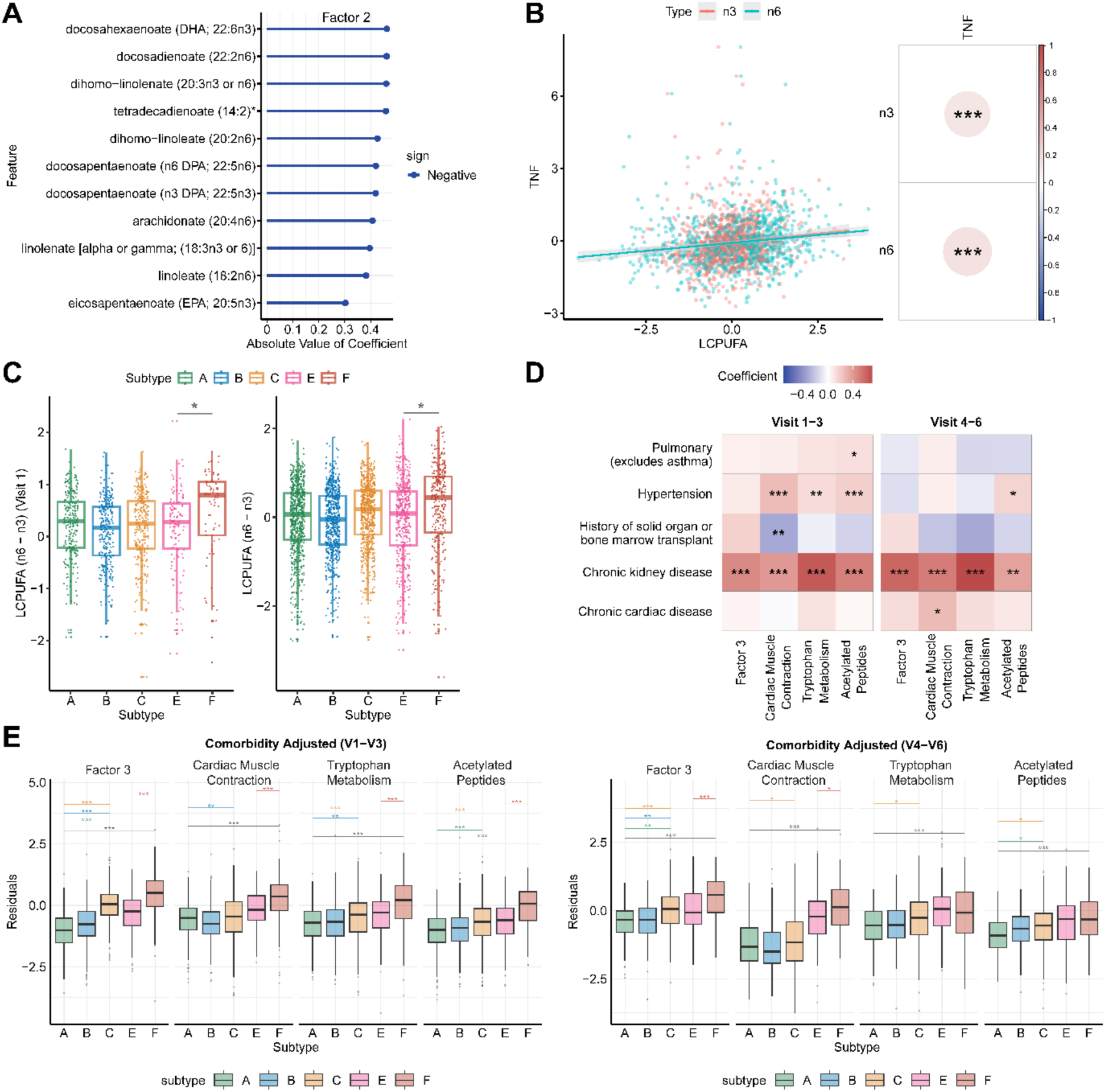
**(A)** Coefficients for long-chain polyunsaturated fatty acids (LCPUFA) (n3 and n6) enriched in Factor 2. **(B)** Scatter plot of TNF against n3 FAs and n6 FAs at visit 1, whose association was test after controlling for age and sex as fixed effects, and enrollment site as random effects. **(C)** Boxplot showing the differences between n6 FAs and n3 FAs across subtypes. The left panel only includes participants at visit 1, while the right panel includes all participants across 6 visits. Significance in the differences between SubF and SubE was evaluated using linear regression after controlling for age and sex, with the right panel further controlled for admission date, and participant ID and enrollment site as random effects. **(D)** Heatmap of most significant comorbidities against Factor 3, along with its associated pathways. Coefficients and significance were derived from the same regression model used in the comorbidity-adjusted analysis in (E). **(E)** Boxplot showing Factor 3 and its associated pathways across subtypes during the early visits and late visits separately, after adjusting for admission date, age, and sex and the participant ID and enrollment site as random effects in the mixed-effect modeling. (*adj.p<=0.05, **adj.p<=0.01, ***adj.p<=0.001).

**Extended Data Figure 6:**
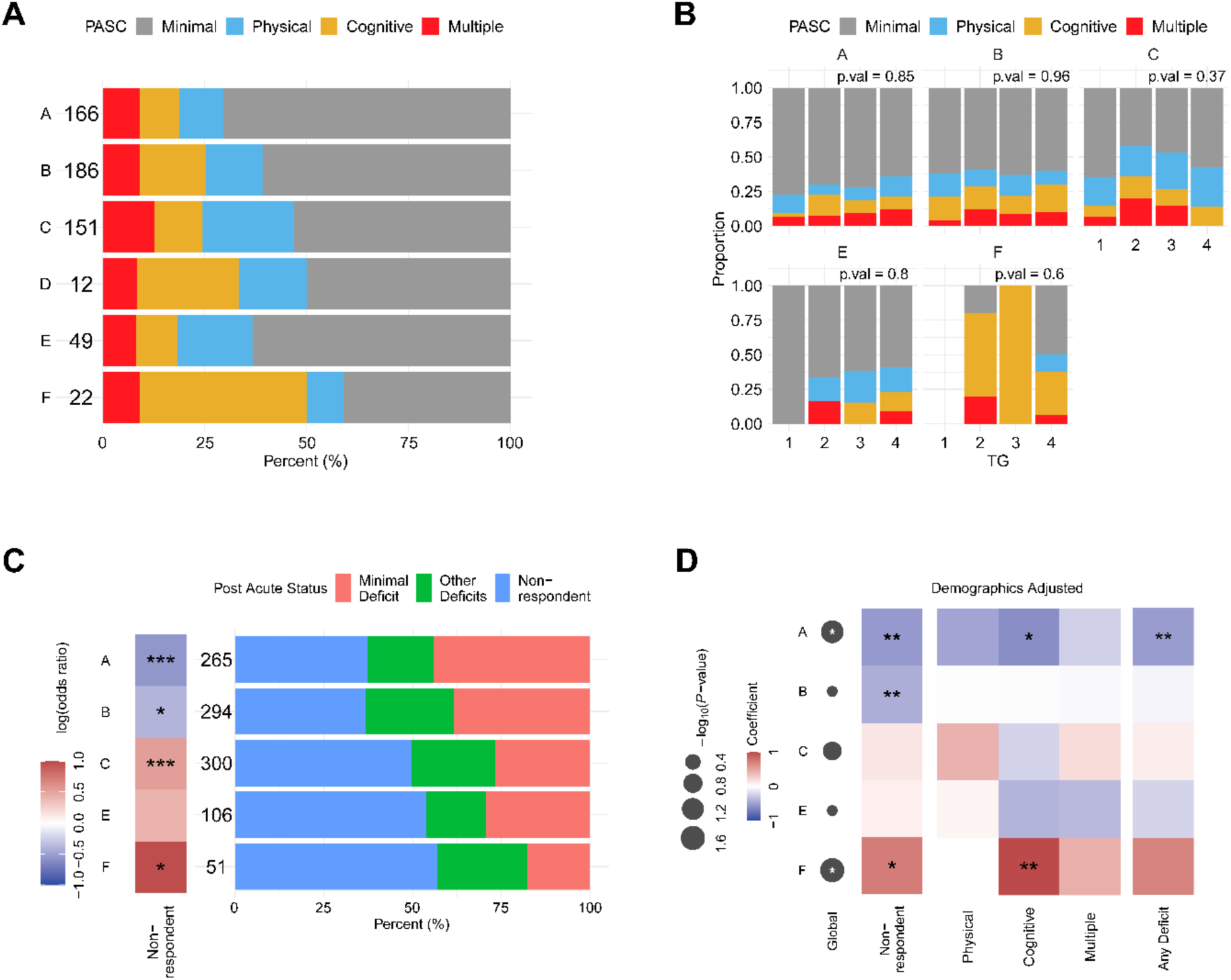
**(A)** Boxplot showing the empirical PASC group proportions across original 6 subtypes including SubD. **(B)** Comparisons of PASC patterns within each subtype using Chi-squared test. **(C)** Comparisons of post-acute status across subtypes using Fisher exact test. Entries in the left heatmap show if non-response or any deficit groups are enriched or not relative to minimal deficit in a given subtype, with colors indicating log odds ratio and significance annotated by asterisk. **(D)** Comparisons of PASC and non-response across subtypes adjusting for demographics. The leftmost row legend shows if the PASC groups were differentially enriched in a given subtype compared to other subtypes after adjusting for age and sex (Global p.val). Entries in the heatmap show if non-response, any deficit, or each deficit PASC group (Physical, Cognitive, and Multiple) is enriched or not relative to the minimal deficit in a given subtype after adjusting for age and sex using logistic regression. (*p<=0.05, **p<=0.01, ***p<=0.001).

**Extended Data Figure 7:**
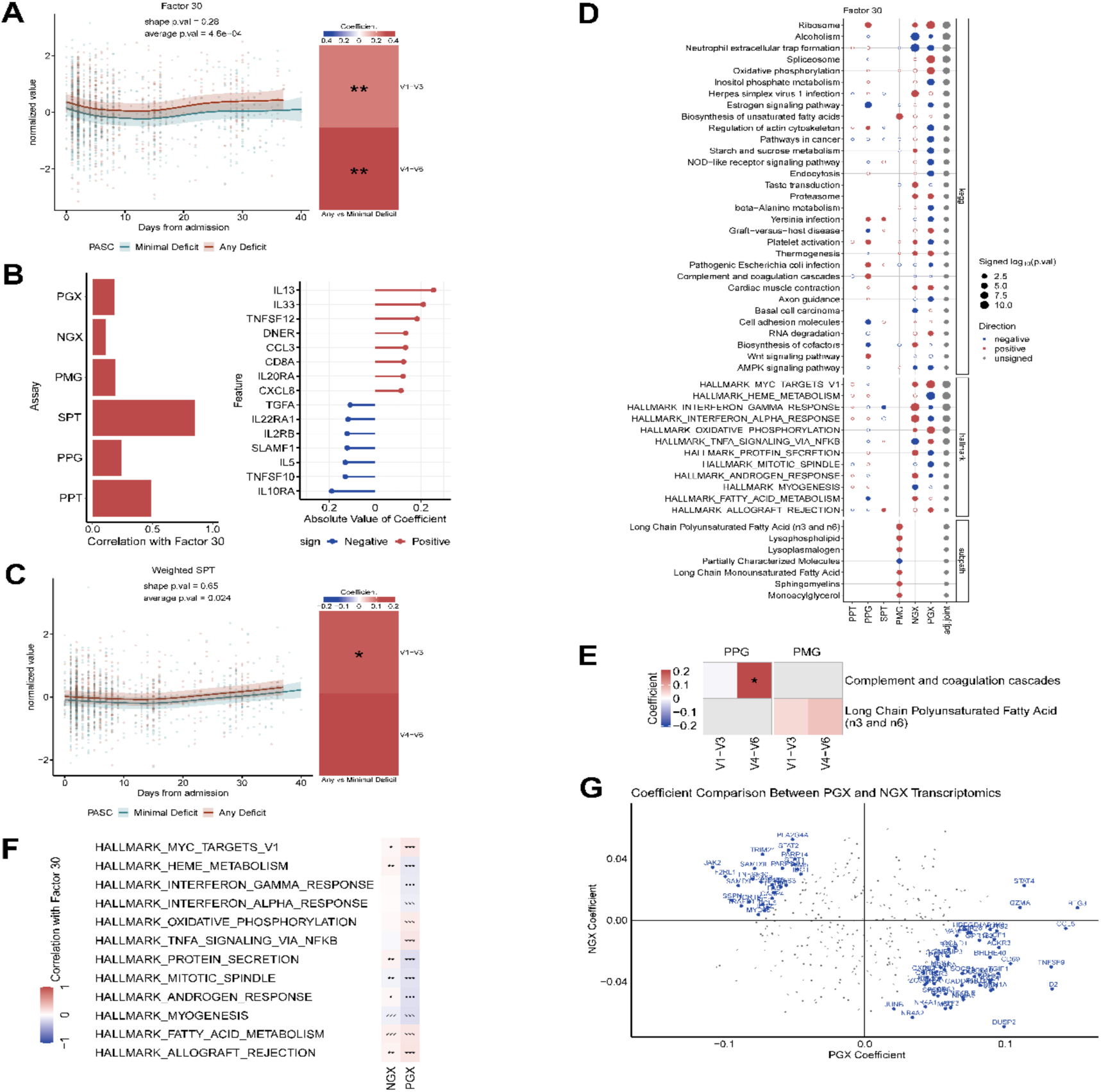
**(A)** Longitudinal trajectories of Factor 30 (left), colored by deficit group with shaded region denoting 95% confidence interval, and heatmaps of comparisons in Factor 30 for visits 1-3 and visits 4-6 among deficit groups (right), colored by regression coefficients. **(B)** Assay correlation with Factor 30 and coefficients of top proteins in SPT. The left barplot shows the correlation between Factor 30 and 6 aggregated assays, weighted by projection coefficients. Aggregated PGX and NGX features showed much weaker associations (COR < 0.2) compared to aggregated SPT features (COR=0.85). The right panel shows the coefficients of top proteins in SPT (absolute coefficient > 0.1). **(C)** Longitudinal trajectories of weighted SPT colored by deficit group, heatmap comparing weighted SPT for visits 1-3 and visits 4-6 among deficit groups. **(D)** Pathway Enrichment of Factor 30 (adj.joint < 0.05). **(E)** Comparison of complement and coagulation in PPG (p.val = 6.4E-1 and 6.3E-2 for early and later visits respectively), and long-chain polyunsaturated fatty acid in PMG (p.val = 2.8E-1 and 7.1E-2 for early and later visits respectively), between any deficit and minimal deficit groups, with colors representing regression coefficients. The top 5 leading-edge proteins in complement and coagulation in Factor 30 include FGG, FGB, C9, C1R, and C1S, which implied pro-coagulation and complement activation (*p<=0.05, **p<=0.01, ***p<=0.001). **(F)** Hallmark pathway correlation with Factor 30. The heatmap shows the correlation between Factor 30 and hallmark pathways in NGX and PGX respectively (*adj.p<=0.05, **adj.p<=0.01, ***adj.p<=0.001). **(G)** Coefficient comparison between PGX and NGX transcriptomics. All genes from the interferon response pathways and TNFA signaling via NFKB pathway, shared by the filtered PBMC and nasal transcriptomics (Supplementary Materials), showed significant negative associations (COR= -0.2, p.val = 1.7E-4).

**Extended Data Figure 8:**
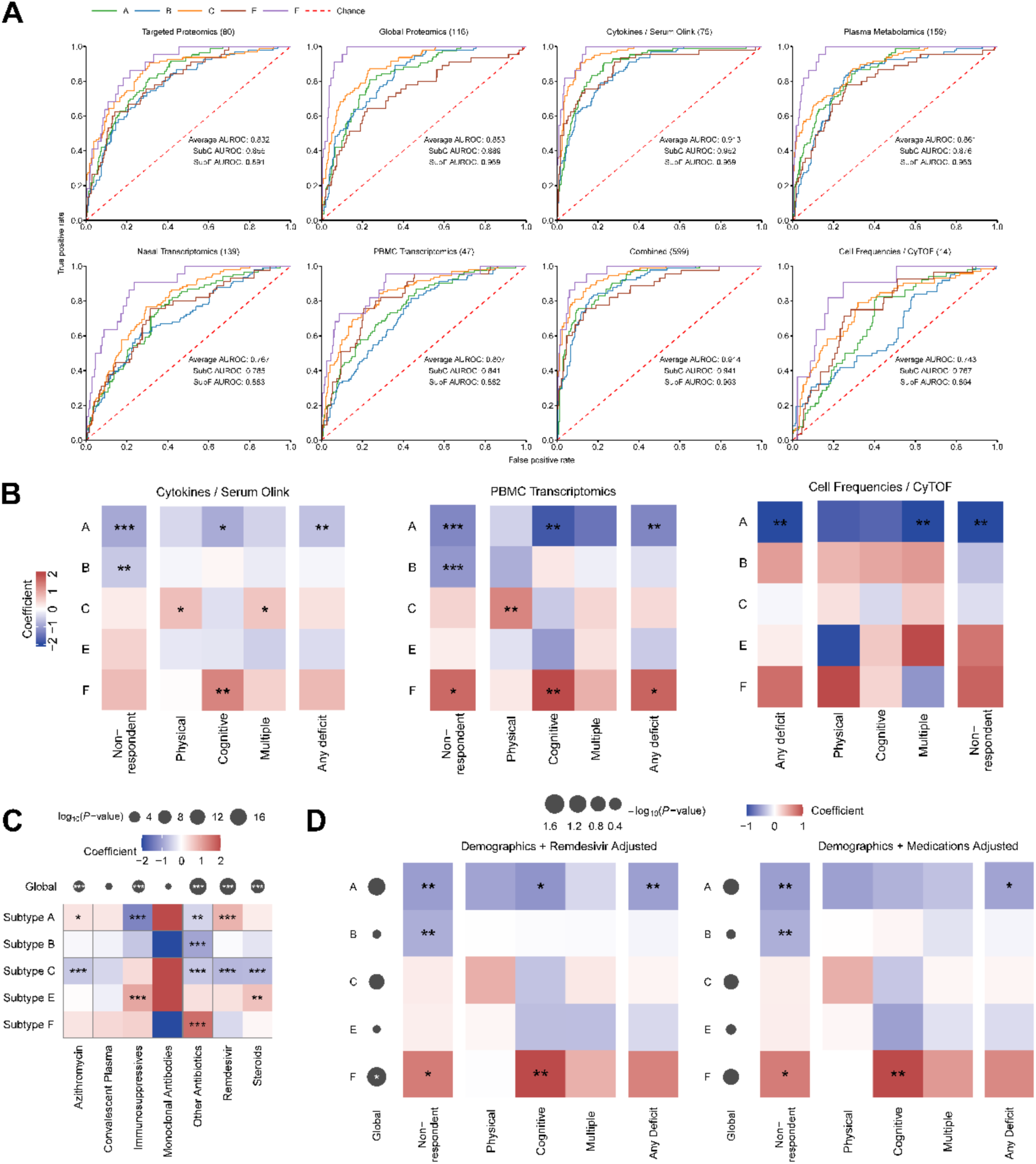
**(A)** AUROC curve for subtype prediction using selected sparse markers from targeted proteomics, global proteomics, serum olink, plasma metabolomics, nasal transcriptomics, and PBMC transcriptomics. The markers were selected using a generalized LASSO linear regression model and subsequently fitted with a radial SVM. **(B)** Heatmap displaying the results of logistic regression on the associations between predicted subtype probabilities from (A) and PASC. (*p<=0.05, **p<=0.01, ***p<=0.001). **(C)** Heatmap of coefficients and BH adjusted p-values comparing medication treatments across subtypes. Each subtype was compared against all others using logistic regressions. The upper row legend shows the Chi-squared test p-values. (*adj.p<=0.05, **adj.p<=0.01, ***adj.p<=0.001). **(D)** Comparisons of PASC and non-response across subtypes, adjusting for age, sex, and remdesivir, as well as for age, sex, and 5 significant medications from (C). (*p<=0.05, **p<=0.01, ***p<=0.001).

